# Sex specific emergence of trisomic *Dyrk1a*-related skeletal phenotypes in the development of a Down syndrome mouse model

**DOI:** 10.1101/2024.05.24.595804

**Authors:** Jonathan M. LaCombe, Kourtney Sloan, Jared R. Thomas, Matthew P. Blackwell, Isabella Crawford, Joseph M. Wallace, Randall J. Roper

## Abstract

Skeletal insufficiency affects all individuals with Down syndrome (DS) or Trisomy 21 (Ts21) and may alter bone strength throughout development due to a reduced period of bone formation and early attainment of peak bone mass compared to typically developing individuals. Appendicular skeletal deficits also appear in males before females with DS. In femurs of male Ts65Dn DS model mice, cortical deficits were pronounced throughout development, but trabecular deficits and *Dyrk1a* overexpression were transitory until postnatal day (P) 30 when there were persistent trabecular and cortical deficits and *Dyrk1a* was trending overexpression. Correction of DS-related skeletal deficits by a purported DYRK1A inhibitor or through genetic means beginning at P21 was not effective at P30, but germline normalization of *Dyrk1a* improved male bone structure by P36. Trabecular and cortical deficits in female Ts65Dn mice were evident at P30 but subsided by P36, typifying periodic developmental skeletal normalizations that progressed to more prominent bone deficiencies. Sex-dependent differences in skeletal deficits with a delayed impact of trisomic *Dyrk1a* are important to find temporally specific treatment periods for bone and other phenotypes associated with Ts21.

**Summary Statement:** Analyzing developing bone and gene expression in Ts65Dn Down syndrome model mice revealed timepoints during development when trisomic *Dyrk1a* overexpression linked to appendicular skeletal abnormalities. *Dyrk1a* was not always overexpressed.

## Introduction

All individuals with Trisomy 21 (Ts21) have skeletal abnormalities and are at risk for osteopenia and osteoporosis. Different from typically developing individuals, both males and females with Down syndrome (DS) exhibit deficits in bone structure and bone accrual during adolescence, and males with DS decline in skeletal parameters earlier than females with DS. A reduced period of bone accrual may also make individuals with DS more susceptible to bone breakage during adolescence (de Moraes et al., 2008). People with DS attain peak bone mass 5-10 years earlier than the general population and experience bone loss sooner, and at a higher rate, than the general population (Carfi et al., 2017, Costa et al., 2018). Growth velocity is reduced in children with DS and skeletal age is delayed compared to chronological age. We have reported that 27% of individuals with DS reported suffering a fracture or broken bone, and most fractures occurred in individuals under 20 years of age (though most individuals that responded were <20 years old) (LaCombe and Roper, 2020). Maximal height in people with DS is reached around age 15, which is precocious compared to the general population (de Moraes et al., 2008, Angelopoulou et al., 1999, Myrelid et al., 2002).

In individuals without DS, age-related trabecular bone loss begins around age 30 and cortical bone loss begins around age 50 (Almeida, 2012, Khosla, 2013, Manolagas et al., 2013). Recent comprehensive studies of adults with DS with large sample sizes have shown significant differences in bone mineral density (BMD) with individuals with DS beginning in their second and third decades of life. Males with DS begin losing BMD in the femur much earlier than females with DS (30 as compared to 40 years of age, respectively), suggesting a protective effect of female sex in terms of maintaining BMD (Carfi et al., 2017, Costa et al., 2017, Costa et al., 2018, Tang et al., 2019). Additionally, because the average life expectancy of individuals with DS has increased to more than 60 years of age (Baird and Sadovnick, 1989, Bittles and Glasson, 2004, Weijerman and de Winter, 2010), more individuals with DS will likely suffer from osteoporosis and fractures in older years. Therefore, there is a critical need to address the etiology, including the impact of trisomic genes, of bone deficiencies and osteoporosis in males and females with DS.

The Ts(17^16^)65Dn (Ts65Dn) mice are the most well-studied mouse model of DS and exhibit many DS-related phenotypes, including skeletal abnormalities (Blazek et al., 2015a, Blazek et al., 2011, Blazek et al., 2015b, Reeves et al., 1995, Thomas et al., 2021). These phenotypes are attributed to the presence of triplicated genes on a freely segregating minichromosome composed of the distal arm of mouse chromosome 16 (Mmu16) attached to the centromeric region of Mmu17. In addition to increased gene dosage of ∼100 genes orthologous to human chromosome 21 (Hsa21), ∼35 protein coding genes are also triplicated in the centromeric region of Mmu17 that are not homologous to Hsa21 (Duchon et al., 2011, Reinholdt et al., 2011). Most studies on Ts65Dn mice have been limited to males, primarily due to their subfertile nature, which requires the use of females for colony maintenance (Roper et al., 2006, Moore et al., 2010). Male Ts65Dn mice have femoral skeletal deficiencies at 6 weeks (a time of bone formation roughly equivalent to the skeletal age of humans under the age of 20) and 16 weeks (a time of skeletal maturity in mice similar to humans between 20 and 30 years of age) (Blazek et al., 2015a, Blazek et al., 2011). Histological assessment of appendicular trabecular bone in male Ts65Dn mice at 6 weeks showed a reduced bone formation rate (BFR), mineral apposition rate (MAR) and mineralizing surface, and increased osteoclast number (Blazek et al., 2015a). At 3 months (∼12 weeks), tibial BFR, percent osteoblast surface/bone surface, percent osteoclast/bone surface, and osteoclast number were reduced in Ts65Dn male mice (Fowler et al., 2012). Bone formation marker P1NP was decreased significantly in Ts65Dn as compared to euploid mice at 24 months and humans with DS at 19-51 years as compared to control individuals, while bone resorption marker TRAP 5b was decreased in Ts65Dn mice at 24 months; however, the CTx bone resorption marker was not significantly decreased in humans with DS aged 19-51 years (Fowler et al., 2012, McKelvey et al., 2013). Female Ts65Dn mice at 6 weeks have also been shown to have trabecular deficits, including lower BMD and increased trabecular separation (Tb.Sp), and cortical deficits, including smaller total cortical cross-sectional area (Tt.Ar), periosteal perimeter (Ps.Pm), and endocortical perimeter (Ec.Pm),compared to female euploid mice (Thomas et al., 2021).

It has been hypothesized that *Dyrk1a*, a gene found in three copies in humans with DS and Ts65Dn mice, significantly contributes to many DS phenotypes including skeletal malformations (Arron et al., 2006, Duchon and Herault, 2016a, Sloan et al., 2023). DYRK1A is a serine-threonine kinase that regulates many downstream proteins and transcription factors including Cyclin D1 and NFAT (Arron et al., 2006, Branchi et al., 2004, Guedj et al., 2012, Park et al., 2009, Atas-Ozcan et al., 2021). Male and female *Dyrk1a* transgenic mice (increased copy number of just *Dyrk1a*) exhibited significantly reduced bone mass including decreased bone volume fraction and reduced trabecular skeletal parameters (Lee et al., 2009). The deficient skeletal phenotype of *Dyrk1a* transgenic mice was characterized by osteoblast deficiencies that resulted in the low bone mass phenotype (Lee et al., 2009). Returning *Dyrk1a* to two functional copies from conception in otherwise trisomic Ts65Dn mice (Ts65Dn,*Dyrk1a*^+/-^) rescued femoral trabecular skeletal parameters in male 6-week-old Ts65Dn mice to euploid levels, improved cortical cross sectional area, and normalized femoral trabecular and cortical MAR and BFR (Blazek et al., 2015a). Yet, normalization of *Dyrk1a* in otherwise trisomic Ts65Dn mice did not have a significant corrective role in developing skeletal abnormalities in at Embryonic day [E}17.5 (Blazek et al., 2015b). These results suggested a time dependent role of trisomic *Dyrk1a* in Ts65Dn mice and possibly individuals with DS.

Several reports describe potential DYRK1A inhibitors and their possible use for correcting DS-related cognitive deficits (Duchon and Herault, 2016b, De la Torre and Dierssen, 2012, Becker et al., 2014). Treatments using supplements containing Epigallocatechin-3-gallate (EGCG)—a putative DYRK1A inhibitor—have been reported to improve cognitive performance in DS mouse models and in some measures in clinical trials (De la Torre et al., 2016, De la Torre et al., 2014, Pons-Espinal et al., 2013, Souchet et al., 2019). In contrast, no significant beneficial effects on cognitive function were found in DS model mice using pure EGCG treatments (Stringer et al., 2015, Stringer et al., 2017a, Goodlett et al., 2020), and evidence shows that high concentrations and prolonged treatment with EGCG harms skeletal structure (Jamal et al., 2022). CX-4945 is a repurposed anti-cancer drug that displays a higher affinity for DYRK1A than harmine and INDY inhibitors (Kim et al., 2016) and was shown to be effective against hematological cancers and reducing osteoclast activity while increasing osteoblast activity, suggesting an overall inhibitory effect on the hematopoietic progenitor lineage and a stimulatory effect on the mesenchymal lineage (Son et al., 2013, Chon et al., 2015). Because of its high affinity for DYRK1A and its impact on osteoblasts and osteoclasts, CX-4945 may be an excellent candidate to treat skeletal abnormalities associated with DS, especially at times when *Dyrk1a* is overexpressed.

We hypothesized that *Dyrk1a*-related appendicular skeletal phenotypes emerge during development between E17.5 and P42 (6 weeks) in Ts65Dn mice and overexpression of *Dyrk1a* during a critical developmental window in the femoral compartment of Ts65Dn mice dysregulates molecular mechanisms that cause aberrant skeletal phenotypes. Like humans with DS, we further posited that there would be a sexual dimorphism in skeletal deficiencies found in Ts65Dn mice. Additionally, we hypothesized this altered expression of *Dyrk1a* may be temporally corrected by genetic and therapeutic means to improve skeletal phenotypes associated with DS in mouse models. Identifying the age that a *Dyrk1a*-related appendicular skeletal phenotype emerges is critical to determine which genetic, molecular, and cellular processes are dysregulated to cause persistent phenotypic changes into adulthood.

## Methods

### Animal Models

Ts(17^16^)65Dn (Ts65Dn) females (Stock 001924), (∼50% C57BL/6 and ∼50% C3H/HeJ advanced intercross background) and B6C3F1 males (Stock 100010) were obtained from the Jackson Laboratory. New Ts65Dn females and B6C3F1 males from the Jackson Laboratory were added to the colony approximately every 6 months to reduce strain variability. *Dyrk1a* heterozygous mutant mice (*Dyrk1a*^+/−^) were obtained from Dr. Mariona Arbones (Fotaki et al., 2002, Fotaki et al., 2004) and were subsequently backcrossed to B6C3F1 mice >10 generations to parallel the genetic background of Ts65Dn mice. B6*.Dyrk1a^tm1Jdc^* (*Dyrk1a^fl/fl^*) mice containing *lox*P sites flanking *Dyrk1a* exons 5 and 6 were obtained from Dr. Jon Crispino (Thompson et al., 2015) and bred to C3H/HeJ mice (Jackson laboratory stock number 000659), resulting in B6C3F1.*Dyrk1a^fl/wt^* offspring, containing *lox*P insertions on one *Dyrk1a* allele. These heterozygous offspring were intercrossed to produce homozygous B6C3.*Dyrk1a^fl/fl^* mice on a similar B6C3 advanced intercross genetic background as the Ts65Dn model. Unless otherwise noted, trisomic and euploid (control) offspring from Ts65Dn × B6C3.*Dyrk1a^fl/fl^*matings were used in these experiments.

B6N.FVB(Cg)-Tg(CAG-rtTA3)4288Slowe/J (Jackson Laboratory Stock Number 016532) reverse tetracycline transactivator (rtTA) mice and B6.Cg-Tg(tetO-cre)1Jaw/J (Jackson Laboratory Stock Number 006234) mice were first intercrossed with C3H/HeJ mice to produce founder strains on a 50% B6 and 50% C3H background similar to Ts65Dn. The B6C3F1 founder strains rtTa and tetO-cre were then crossed and male progeny from that cross that were positive for both rtTA and tetO-cre and were mated to Ts65Dn, *Dyrk1a*^fl/wt^ females to generate test mice that would experience the loss of a floxed *Dyrk1a* allele after treatment with doxycycline, by inducing activation of the tetracycline-responsive promotor element and excision of exon 5-6 of *Dyrk1a* by Cre on one Mmu16.

Mice were group-housed with mixed genotypes according to sex in a 12:12 light:dark cycle with white light off between 19:00-07:00. Femurs obtained from mice euthanized at specified time points were wrapped in gauze, soaked in phosphate buffered saline (PBS), immersed quickly in liquid nitrogen, and stored at -80°C until ready to use. Right femurs were subjected to microcomputed tomography (μCT) analysis and left femurs were reserved for gene expression analysis. Experiments with animals were carried out in accordance with the NIH Guide for the Care and Use of Laboratory Animals and received prior approval from the IUPUI School of Science IACUC (SC298R and SC338R).

### Genotyping

Ts65Dn mice were genotyped to determine the presence of a freely segregating chromosome by amplifying the Mmu16/Mmu17 breakpoint using primers GTGGCAAGAGACTCAAATTCAAC and TGGCTTATTATTATCAGGGCATTT (Reinholdt et al., 2011). *Dyrk1a*^fl/wt^ mice were genotyped by PCR as described by The Jackson Laboratory protocol using primers TACCTGGAGAAGAGGGCAAG and GGCATAACTTGCATACAGTGG. Experiments using the *Dyrk1a* heterozygous mutant model confirmed the presence of the mutated allele with primers ATTCGCAGCGCATCGCCTTCTATCGCC and CGTGATGAGCCCTTACCTATG using a previously described protocol (Fotaki et al., 2002).

Presence of the rtTA allele was determined by PCR using primers CTGCTGTCCATTCCTTATTC, CGAAACTCTGGTTGACATG, and TGCCTATCATGTTGTCAAA to produce the carrier 330bp and WT 363bp bands (Paterson et al., 2015, Takiguchi et al., 2013). Presence of the tetO-cre allele in test mice was determined by PCR using Cre primers ATTCTCCCACCGTCAGTACG and CGTTTTCTGAGCATACCTGGA (Chai et al., 2000, Miwa et al., 2009) and internal positive control primers CAAATGTTGCTTGTCTGGTG and GTCAGTCGAGTGCACAGTTT (Deitz and Roper, 2011) to produce the carrier 475bp and WT 200bp band when resolved on a 1.5% agarose gel.

### Cre Activation

Doxycycline was administered in chow (ENVIGO, Teklad Custom Diet TD.120769, 998.975g/kg 2018 Teklad Global 18% Protein Rodent Diet, 0.625g/kg Doxycycline hyclate, 0.4g/kg Blue Food Color), delivering an average daily dose of 2-3mg doxycycline per 4-5g chow consumed per day. Doxycycline chow was introduced at the time of weaning, postnatal day 21 (P21). Truncation of *Dyrk1a* was verified at P30 by PCR using tail DNA. Excision of the floxed region spanning exons 5 and 6 on one allele of *Dyrk1a* in Ts65Dn, *Dyrk1a*^fl/wt^, tetO-cre^+^, rtTA^+/-^ mice that had received doxycycline feed was verified with PCR using primers ACCTGGAGAAGAGGGCAAGA and GCCACTGTGTGAGGAGTCTT (Thomas et al., 2021).

### Micro Computed Tomography (μCT) and Analysis

Scans and analysis were performed on right femurs as described (Sloan et al., 2023) using a SkyScan 1172 high-resolution μCT system (Bruker, Kontich, Belgium). Flat field corrections occurred prior to scanning. Hydroxyapatite phantoms (0.25 and 0.75g/cm^3^ CaHA) were scanned once per week of scanning. Scanning parameters are as follows: 60kV, 12µm resolution, 885ms integration time, 0.7° angular increment, frame averaging of 4. The entire length of the femurs from P12, P15, P18, P24, and P27 were scanned, but due to size constraints, samples from P30 and P42 were scanned from the distal condyles of the femur to at least the third trochanter. Trabecular microarchitecture was analyzed by identifying a trabecular region of interest (ROI) beginning at the proximal end of the distal growth plate and extending proximally based on 10% of the bone length from the proximal end of the distal growth plate to the widest region of the third trochanter using SkyScan CT Analyzer (CTAn) and a custom MATLAB code to exclude the outer cortical bone, using lower greyscale threshold of 45 (P12 mice) or 55 (all other ages) and upper greyscale threshold of 255, as described in previous publications (Goodlett et al., 2020, Stringer et al., 2017a, Thomas et al., 2020, Sloan et al., 2023). The cortical ROI was defined as a region of seven transverse slices at 60% of the overall length of the bone from the proximal end of the distal growth plate and cortical geometry was measured as previously described using MATLAB (Goodlett et al., 2020, Stringer et al., 2017a, Thomas et al., 2020, Sloan et al., 2023). Skeletal geometry measures are expressed as standardized nomenclature, symbols, and units for bone histomorphometry according to ASBMR guidelines (Dempster et al., 2013).

### Femoral RNA Isolation

RNA was isolated from the cortical midshaft left femora using Trizol-chloroform as summarized below. After thawing to room temperature (RT), the proximal and distal ends of femora were excised using a razor blade. The cortical midshaft was placed in a modified 200μL pipette tip and located inside a 1.5mL microcentrifuge tube. Femora were centrifuged for 10min at 13,200rpm (Eppendorf® 5702 Centrifuge with F45-24-11 rotor) to remove marrow. Femora were then placed into a pre-chilled mortar containing liquid nitrogen and pulverized into a fine powder. Bone powder was then scraped into a nuclease-free 1.5mL microcentrifuge tube and combined with 50μL Trizol before homogenizing with a motorized hand homogenizer for 60sec. Trizol (150μL) was added and homogenized again. Trizol (300μL) was added, the sample was vortexed, and then centrifuged at 4°C for 10min at 10,000rpm. The supernatant was transferred to another 1.5mL tube and the process was repeated by adding 50μL, 150μL, and 300μL of Trizol to the pellet after intermittent mixing with a hand homogenizer and vortexing. After the supernatant was incubated at RT for 5min, chloroform was added to the supernatant and shaken for 15sec. The aqueous and organic layer were allowed to separate by incubation at RT for 5min. The samples were then centrifuged at 4°C for 10min at 10,000rpm after which the aqueous layer (top) was transferred to a new tube. RNA was precipitated by adding 800μL 100% isopropanol. After a 5min incubation at RT, samples were centrifuged at 4°C for 15min at 13,200rpm. Supernatant was discarded, 1mL of 75% ethanol (EtOH) was added to each sample, and shaken for 15sec after which samples were centrifuged at 4°C for 15min at 13,200rpm. The supernatant was discarded and centrifuged again at 4°C for 1min at 13,200rpm to consolidate the supernatant. The remaining supernatant was aspirated, and tubes were left open for 5min for EtOH evaporation. RNA was resuspended with 20μL of HPLC grade water, concentration was quantified by ThermoScientific NanoDrop 2000, and RNA stored at -80°C for future use.

### cDNA Conversion and Gene Expression Analysis by qPCR

Isolated and quantified RNA samples were converted to cDNA with a final volume of 20μL using TaqMan™ Reverse Transcription Reagents following the recommended protocol, except for doubling the time of the elongation step. The resulting product was diluted 1:5 with sterile Milli-Q™ (MilliporeSigma) water and stored at -20°C until analysis. RT-qPCR was performed using an Applied Biosystems 7500 Real-Time PCR System on 96 well plates.

Reagents consisted of TaqMan™ Gene Expression Master Mix (Roche Diagnostics), TaqMan control probes for *Rn18s* (Roche Diagnostics, Mm03928990_g1), two probes for different regions of the *Dyrk1a* transcript, the first probe targeting a region spanning *Dyrk1a* exons 5-6 (Mm00432929_m1) and the second targeting *Dyrk1a* exons 10-11 (Mm00432934_m1), *Bglap* (Mm03413826_m1), *Runx2* (Mm00501584_m1), *Rbl2* (Mm01242468_m1), and *Alpl* (Mm00475834_m1).

The ΔRn threshold was set at 0.1. ΔCT=CT*_Dyrk1a_* – CT*_Rn18s_*. Relative gene expression = 2^-ΔCT^=ΔCT_Ts65Dn_ – ΔCT_WT_. Fold change=2^-ΔCT^_WT_/2^-ΔCT^_Ts65Dn_. The normalized reporter (Rn) was defined as the difference between the reporter signal, FAM, and the quencher, ROX. The difference between the Rn of a given cycle and the Rn of the first cycle was defined as ΔRn. The cycle threshold (Ct) of 0.100 was chosen because it was a positive threshold that captured the exponential growth phase for all target genes. The difference between the gene of interest *Dyrk1a* and the housekeeping gene *Rn18s* in each sample was defined as ΔCt. Since *Rn18s* expression is conserved between genotypes, the ΔCt calculation produced a value that would be different between genotypes if *Dyrk1a* was overexpressed. To obtain a relative gene expression, each individual sample measure was normalized using the 2^-ΔCT^ method. To determine the fold change of *Dyrk1a* expression between genotypes, each 2^-ΔCT^ was divided by the mean euploid 2^-ΔCT^ and the average of the relative fold change was used to quantify *Dyrk1a* gene expression. The resulting values can only be positive; overexpression of a gene would be indicated by a value greater than 1, while under-expression of a gene would be indicated by a value between 0 and 1.

### Treatment of CX-4945 in Ts65Dn,Dyrk1a^fl/wt^ Mice

Ts65Dn,*Dyrk1a*^fl/wt^ and control male mice were treated with 75mg/kg/day CX-4945 (Silmitasertib, SelleckChem) as a 10% DMSO/90% PBS suspension delivered via oral gavage. A 250mM solution with 24.5mg CX-4945 and 280μL DMSO was gently heated in a 37°C water bath or heat block. The working suspension was made daily by diluting the stock solution to 25mM with PBS to make a suspension (10% total DMSO) and kept in a 37°C heat block until administered to weaned (P21) to P29 mice. Control animals received a vehicle composed of 10% DMSO and 90% PBS.

### Statistical Analyses

Skeletal parameters from μCT were analyzed using IBM SPSS Statistics (v 29.0.1.0). Normality was assessed using Shapiro-Wilk’s test (ɑ=0.05), and non-normal data was logarithmically transformed and tested again. Homogeneity of variance was assessed using Levene’s test (ɑ =0.05). *P* values generated from analysis of variance (ANOVA) tests and t-tests were adjusted using the Benjamini-Hochberg method of False Rate Discovery (FDR) correction separately for trabecular and cortical parameters. Significance was determined as an adjusted *p*≤0.05. In cases where Levene’s test was significant, indicating violation of homogeneity of variance, Welch’s t-test was used to confirm significance (ɑ=0.05), and a parameter was reported as significant if both the FDR-adjusted ANOVA and Welch’s t-test produced a significant *p* value. For development of Ts65Dn skeletal phenotype from P12-P42 experiment, two-way ANOVAs between genotype, age, and their interaction was performed with Tukey’s (non-significant Levene’s test) or Games-Howell (significant Levene’s test) *post hoc* analysis in cases of a significant age effect (no interactions were detected). Additionally, two-tail t-tests were performed between the two genotypes for each age. For the CX-4945 treatment experiment, two-way ANOVAs between genotype, treatment, and their interaction were performed. Additionally, a repeated-measures ANOVA was performed for body weight, using genotype and treatment as between subject factors and postnatal day as the within subject factor. Sphericity was assessed using Mauchly’s test, and Greenhouse-Geisser test was utilized for within subject effect due to significant Mauchly’s test. Pairwise comparisons were utilized in cases where within subject effect or interactions involving within subject effect were significant. For the temporal reduction of *Dyrk1a* experiment, one-way ANOVAs were performed between the four genotype groups with Tukey’s or Games-Howell *post hoc* analysis when significant. With the P30 and P36 germline reduction of *Dyrk1a* copy number experiments, two-way ANOVAs were performed between genotype, sex, and their interaction with Tukey’s or Games-Howell *post hoc* analysis in cases of a significant genotype effect and/or interaction.

Additionally, one-way ANOVAs were performed between the four genotype groups for each sex with Tukey’s or Games-Howell *post hoc* analysis when significant. Significance for qPCR *Dyrk1a* and other osteogenic gene expression measures was determined by unpaired one-tailed t-tests. Normality was not violated at any age (GraphPad Prism 9.0.1).

## Results

### Development of trabecular deficits in male Ts65Dn DS model mice from P12-P42

We have observed trabecular and cortical skeletal deficits in 6- and 16-week DS mouse models that vary across age (Blazek et al., 2011, Sloan et al., 2023, Thomas et al., 2020, Thomas et al., 2021). To further understand the development of these skeletal deficits, we first examined trabecular and cortical bone development in male Ts65Dn DS model and euploid littermate control mice from postnatal day (P)12 to P42. We detected significant genotype and age effects when trabecular bone measurements were analyzed together from P12-42 (Table 1). When trabecular phenotypes at each age were analyzed independently, we found at P12 and P15, there were no significant differences between trisomic and euploid mice; at P18, Ts65Dn male mice had significantly reduced bone mineral density (BMD) (p=0.034) and bone volume fraction (BV/TV) (p=0.023) and increased trabecular separation (Tb.Sp) (p=0.035) (Fig. 1A-E). Additionally, trabecular deficits were not seen in trisomic male mice at P24 and P27. At P30, trabecular phenotypes characteristic of adult male Ts65Dn mice emerged, evidenced by significant reduction in BMD (p=0.030), BV/TV (p=0.030), and trabecular number (Tb.N) (p=0.033) and an increase in Tb.Sp (p=0.050). Similar to what has been reported at P42 (6 weeks), we found P42 Ts65Dn as compared to euploid mice had significantly reduced BMD (p=0.040), BV/TV (p=0.025), Tb.Th (p=0.023), Tb.N (p=0.043) and significantly increased Tb.Sp (0.036).

**Figure 1:**
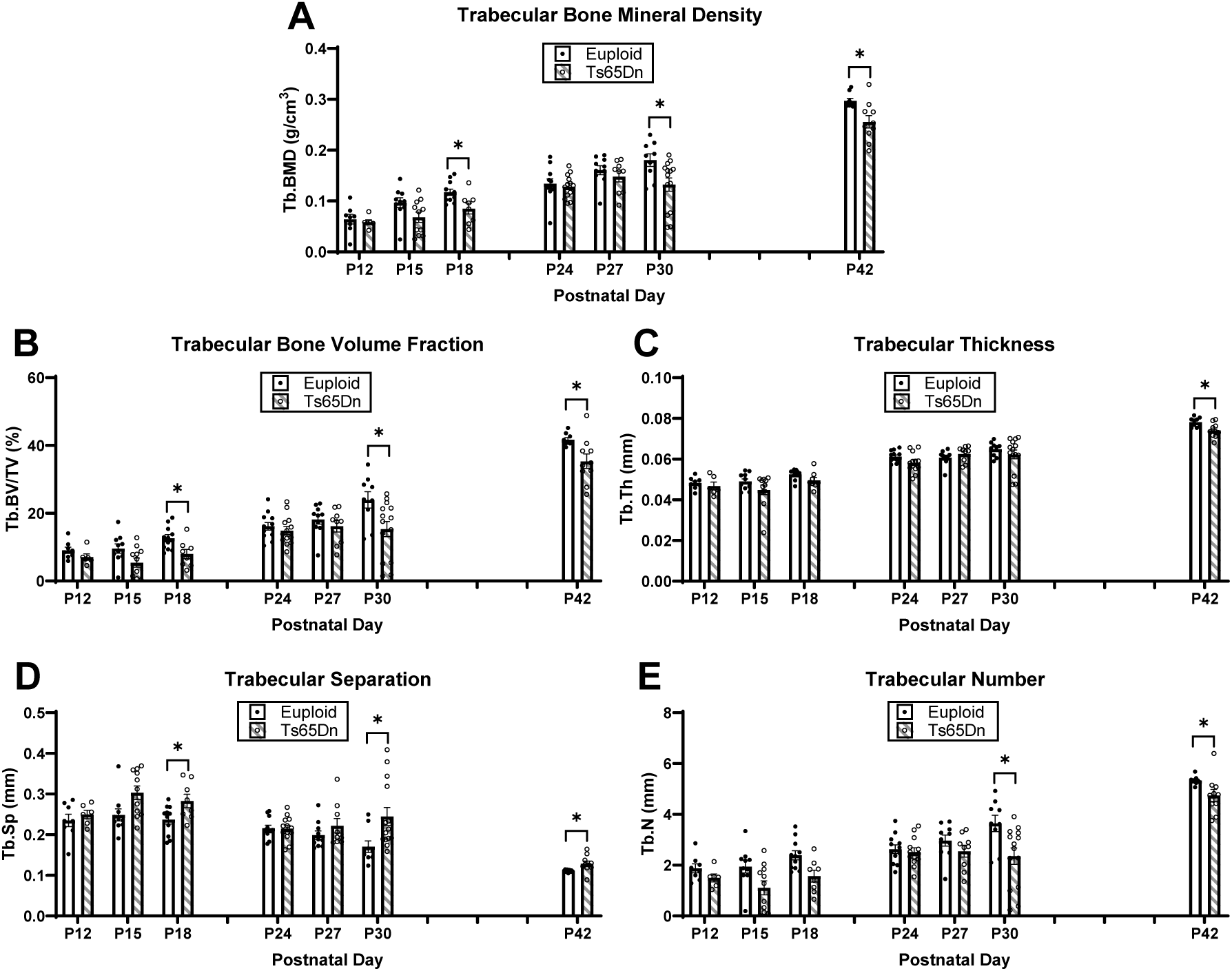
Trabecular analysis of male Ts65Dn femurs from postnatal day (P)12-42. Trabecular parameters are from µCT. Data are mean ± SEM. Significance determined through two-tail t-test with FDR adjustment. (*) indicates p ≤ 0.05, (**) indicates p ≤ 0.01. P12: euploid (n = 8), Ts65Dn (n = 6); P15: euploid (n = 10), Ts65Dn (n = 11); P18: euploid (n = 11), Ts65Dn (n = 8); P24: euploid (n = 12), Ts65Dn (n = 13); P27: euploid (n = 10), Ts65Dn (n = 9); P30: euploid (n = 9), Ts65Dn (n = 14); P42 euploid (n = 9), Ts65Dn (n = 10).

**Table 1:** Significant *p* values from two-way ANOVA (genotype x age) for male P12-P42 Ts65Dn trabecular and cortical parameters. All effect and interaction *p* values reported are adjusted using the Benjamini-Hochberg method. Welch’s t-test was performed to confirm ANOVA results when a significant Levene’s test occurred, indicating unequal variances between groups, and these *p* values are unadjusted. If Welch’s t-test encountered a non-significant result when ANOVA generated a significant result, the non-significant result was reported. ǂ indicates the data was log-transformed for statistical analysis due to non-normal distribution. NS indicates *p* value was not significant. N/A indicates Welch’s t-test was not performed because Levene’s test was not significant. See Figure 1 for group numbers.

|  | BMD | BV/TV | Tb.Th | Tb.Sp | Tb.N |  |  |  |  |
| --- | --- | --- | --- | --- | --- | --- | --- | --- | --- |
| **Genotype Effect** | 0.003 | 0.003 | 0.008 | 0.003 | 0.003 |  |  |  |  |
| Welch's t-test | NS | NS | NS | 0.007 | 0.014 |  |  |  |  |
| **Age Effect** | 0.003 | 0.003 | 0.003 | 0.003 | 0.003 |  |  |  |  |
| Welch's t-test | <0.001 | <0.001 | <0.001 | <0.001 | <0.001 |  |  |  |  |
| **Interaction** | NS | NS | NS | NS | NS |  |  |  |  |
|  | Tt.Ar<sup>†</sup> | Ma.Ar<sup>†</sup> | Ct.Ar<sup>†</sup> | Ct.Th | Ps.Pm | Ec.Pm | I<sub>max</sub><sup>†</sup> | I<sub>min</sub><sup>†</sup> | Ct.TMD |
| **Genotype Effect** | 0.003 | 0.003 | 0.003 | 0.003 | 0.003 | 0.003 | 0.003 | 0.003 | 0.003 |
| Welch's t-test | <0.001 | N/A | 0.004 | NS | <0.001 | N/A | <0.001 | <0.001 | NS |
| **Age Effect** | 0.003 | 0.003 | 0.003 | 0.003 | 0.003 | 0.003 | 0.003 | 0.003 | 0.003 |
| Welch's t-test | <0.001 | N/A | <0.001 | <0.001 | <0.001 | N/A | <0.001 | <0.001 | <0.001 |
| **Interaction** | NS | NS | NS | NS | NS | NS | NS | NS | NS |

### Cortical deficits in male Ts65Dn DS model mice from P12-P42

Significant genotype and age effects were found when cortical bone measures were analyzed from P12-P42 (Table 1). Examining specific cortical phenotypes at each age found that at P12, there were significant cortical deficits in all cortical measures in Ts65Dn as compared to euploid male mice: total cross-sectional area (Tt.Ar) (p=0.011), marrow area (Ma.Ar) (p=0.032), cortical area (Ct.Ar) (p=0.011), cortical thickness (Ct.Th) (p=0.019), periosteal perimeter (Ps.Pm) (p=0.012), endocortical perimeter (Ec.Pm) (p=0.034), maximum moment of inertia (I_max_) (p=0.009), minimum moment of inertia (I_min_) (p=0.009) and cortical tissue mineral density (Ct.TMD) (p=0.009) (Fig. 2A,C-D). At P15, there was a significant reduction in all cortical measures (except Ct.TMD) in Ts65Dn as compared to euploid littermate male mice: Tt.Ar (p=0.005), Ma.Ar, (p=0.005), Ct.Ar (p=0.005), Ct.Th (p=0.028), Ps.Pm (p=0.005), Ec.Pm (p=0.005), I_max_ (p=0.006) and I_min_ (p=0.006). In P18, all cortical measurements (including Ct.TMD) were significantly reduced in trisomic as compared to euploid male mice: Tt.Ar (p=0.002), Ma.Ar (p=0.002), Ct.Ar (p=0.002), Ct.Th (p=0.002), Ps.Pm (p=0.002), Ec.Pm (p=0.002), I_max_ (p=0.002), I_min_ (p=0.002), and Ct.TMD (p=0.002). At P24, only Ct.Ar (p=0.027), Ct.Th (p=0.014), I_min_ (p=0.045), and Ct.TMD (0.008) were significantly reduced in trisomic as compared to control male mice. At P27, cortical defects were again significantly different in Tt.Ar (p=0.005), Ma.Ar (p=0.005), Ps.Pm (p=0.004), Ec.Pm (p=0.004), I_max_ (p=0.023), and I_min_ (p=0.005). At P30, all cortical parameters were significantly reduced in trisomic as compared to euploid male mice: Tt.Ar (p=0.005), Ma.Ar (p=0.004), Ct.Ar (p=0.004), Ct.Th (p=0.004), Ps.Pm (p=0.004), Ec.Pm (p=0.004), I_max_ (p=0.004), I^min^ (p=0.005), and Ct.TMD (p=0.005). At P42, similar to what has been previously reported, all cortical measurements except Ct.Th were significantly reduced in male Ts65Dn as compared to euploid mice: Tt.Ar (p=0.002), Ma.Ar (p=0.002), Ct.Ar (p=0.002), Ps.Pm (p=0.002), Ec.Pm (p=0.002), I_max_ (p=0.002), I_min_ (p-0.002), and Ct.TMD (p=0.005).

**Figure 2:**
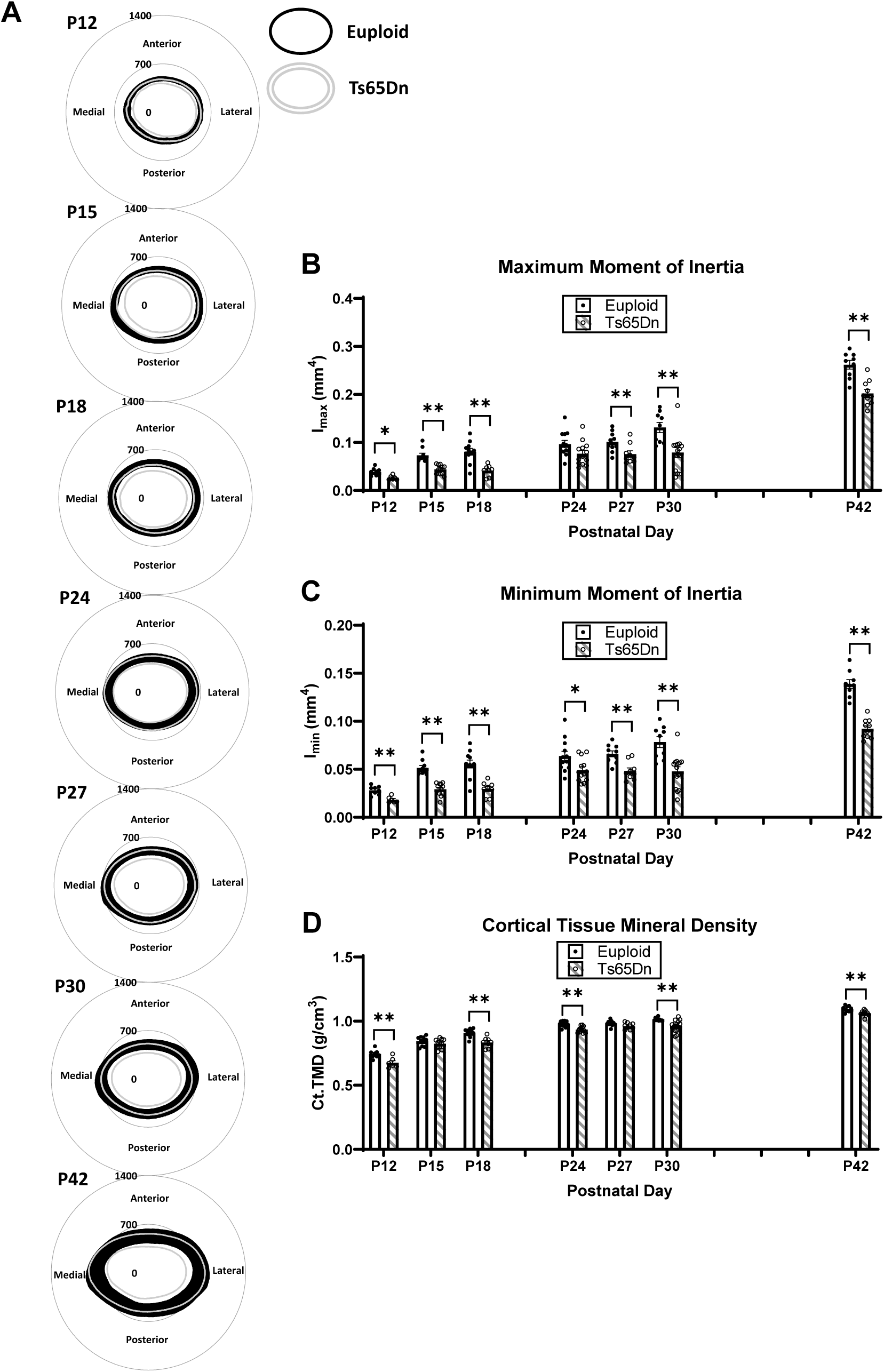
Cortical analysis of male Ts65Dn femurs from postnatal day (P)12-42. (A) Cortical models of euploid (filled in black) and Ts65Dn (grey outline) made by finding the centroid of the bone then calculating inner and outer radius measurements every 0.5 degrees in MATLAB for each cortical slice (total of 7) of each animal, then finding the average for the entire group. Scale is in microns. (B-D) Some cortical parameters from μCT analysis (see Supplemental Figure 1 for remaining parameters). Data are mean ± SEM. Significance determined through two-tail t-test with FDR adjustment. (*) indicates p ≤ 0.05, (**) indicates p ≤ 0.01. See Figure 1 for group numbers.

Trabecular and cortical data were then plotted on an age timeline from P12-P42 and assessed by two-way ANOVA with age and genotype as factors to understand when trabecular and cortical bone growth were occurring. This perspective revealed stagnant periods of growth in euploid and Ts65Dn males with no significant growth (based on lack of age effect) from P12 to P18 and from P24 to P30 for most trabecular parameters (Fig. 3). Stagnant periods of growth in cortical bone were dependent on parameter: Tt.Ar, Ps.Pm, and Ec.Pm did not significantly change from P15 to P30, Ma.Ar did not change from P15 to P42, and Ct.Ar, Ct.Th, I_max_, I_min_, and Ct.TMD had two periods from P15 to P18 and P24 to P30 (Fig. 4). Overall, this indicated two major periods of trabecular bone growth (P18 to P24, then P30 to P42) and three major periods of cortical bone growth (P12 to P15, then P18 to P24, then P30 to P42). These data prompted a deeper investigation into genetic mechanisms during the growth period spanning P12 to P42.

**Figure 3:**
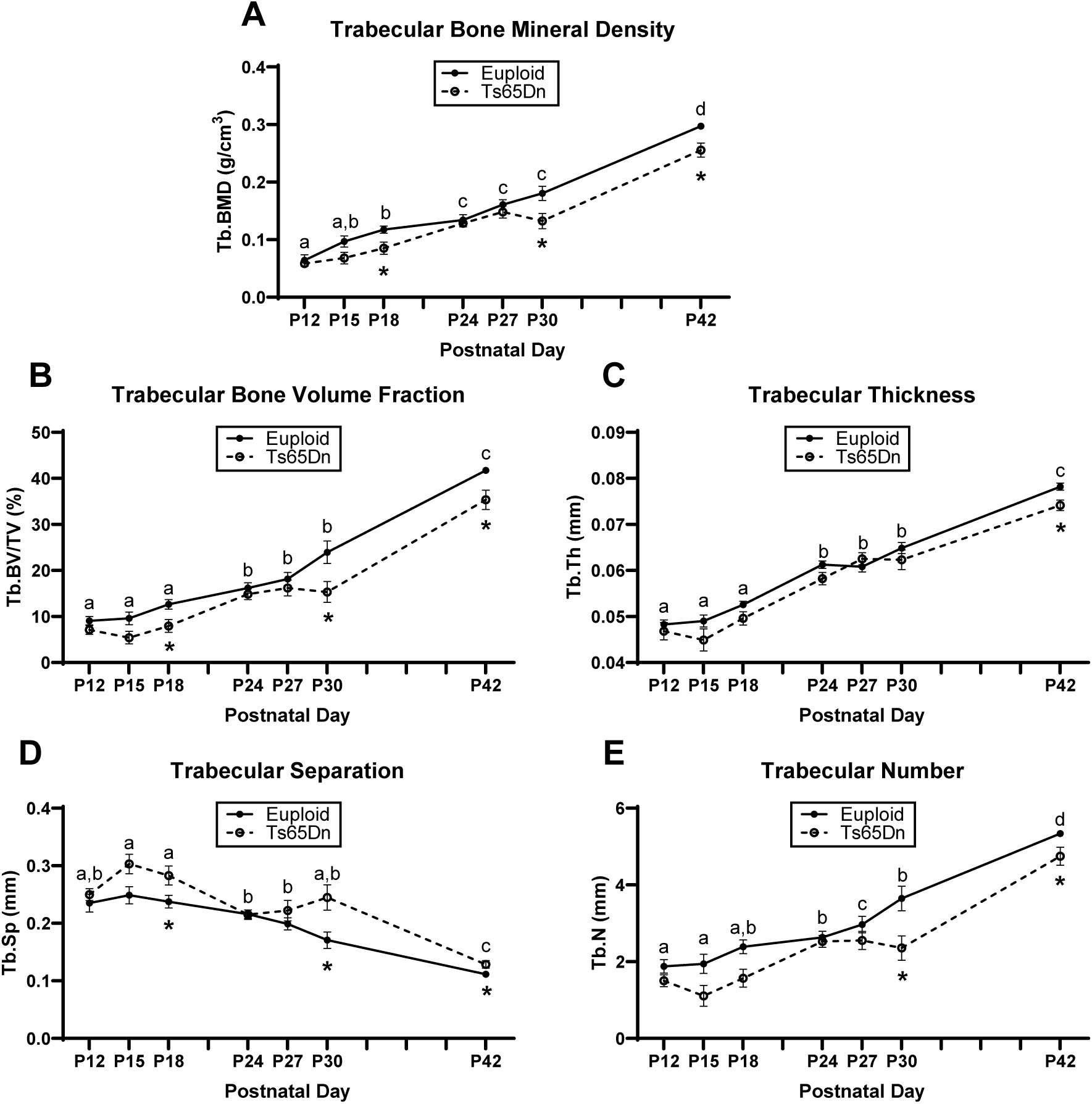
Cross-sectional growth trajectory of trabecular bone in male euploid and Ts65Dn femurs from postnatal day (P)12-42. Data are mean ± SEM. Letters (a,b,c,d) indicate significant differences between ages by two-way ANOVA and Tukey/Games-Howell *post hoc* analysis (age effect); ages with the same letter are not significantly different from one another. (*) indicates adjusted p ≤ 0.05 by two-tail t-test between euploid and Ts65Dn mice at the given age. See Figure 1 for group numbers.

**Figure 4:**
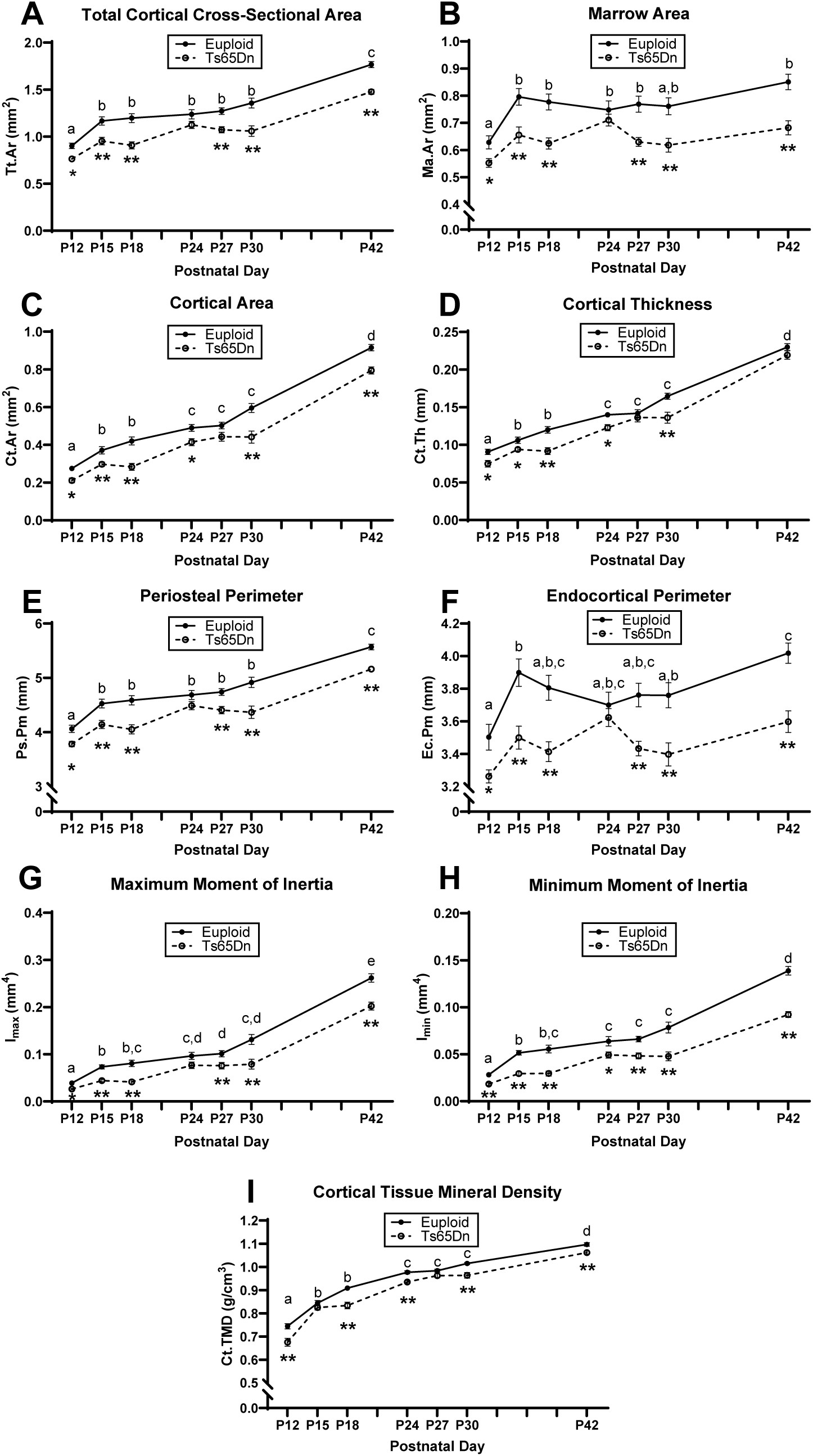
Cross-sectional growth trajectory of cortical bone in male euploid and Ts65Dn femurs from postnatal day (P)12-42. Data are mean ± SEM. Letters (a,b,c,d,e) indicate significant differences between ages by two-way ANOVA and Tukey/Games-Howell *post hoc* analysis (age effect); ages with the same letter are not significantly different from one another. (*) indicates adjusted p ≤ 0.05 by two-tail t-test between euploid and Ts65Dn mice at the given age. (**) indicates adjusted p ≤ 0.01 by two-tail t-test between euploid and Ts65Dn mice at the given age. See Figure 1 for group numbers.

### Relationship between Dyrk1a expression and bone development in male Ts65Dn mice

We have previously shown that three copies of *Dyrk1a* is an important factor in causing trabecular and cortical cross-sectional area deficits in 6-week-old Ts65Dn male mice (Blazek et al., 2015a). To test the hypothesis that *Dyrk1a* overexpression during postnatal development in male Ts65Dn mice is linked to the emergence of appendicular skeletal abnormalities, RNA was isolated from the midshafts of left femurs obtained from P12, 15, 18, 24, 27, and 30 Ts65Dn and euploid male mice. At P12, male trisomic *Dyrk1a* was significantly overexpressed with a fold change of 2.053 (p=0.041) (Fig. 5). The male trisomic *Dyrk1a* fold change at P15 was 1.037 (p=0.417). *Dyrk1a* was significantly overexpressed at P18; the fold change for male Ts65Dn mice was 2.03 (p=0.011). At P24, the male trisomic *Dyrk1a* fold change value was determined to be 0.76 (p=0.238). The *Dyrk1a* fold change at P27 for Ts65Dn male mice was 1.61 (p=0.072). The *Dyrk1a* fold change for Ts65Dn male mice at P30 was 1.81 (p=0.0598).

**Figure 5:**
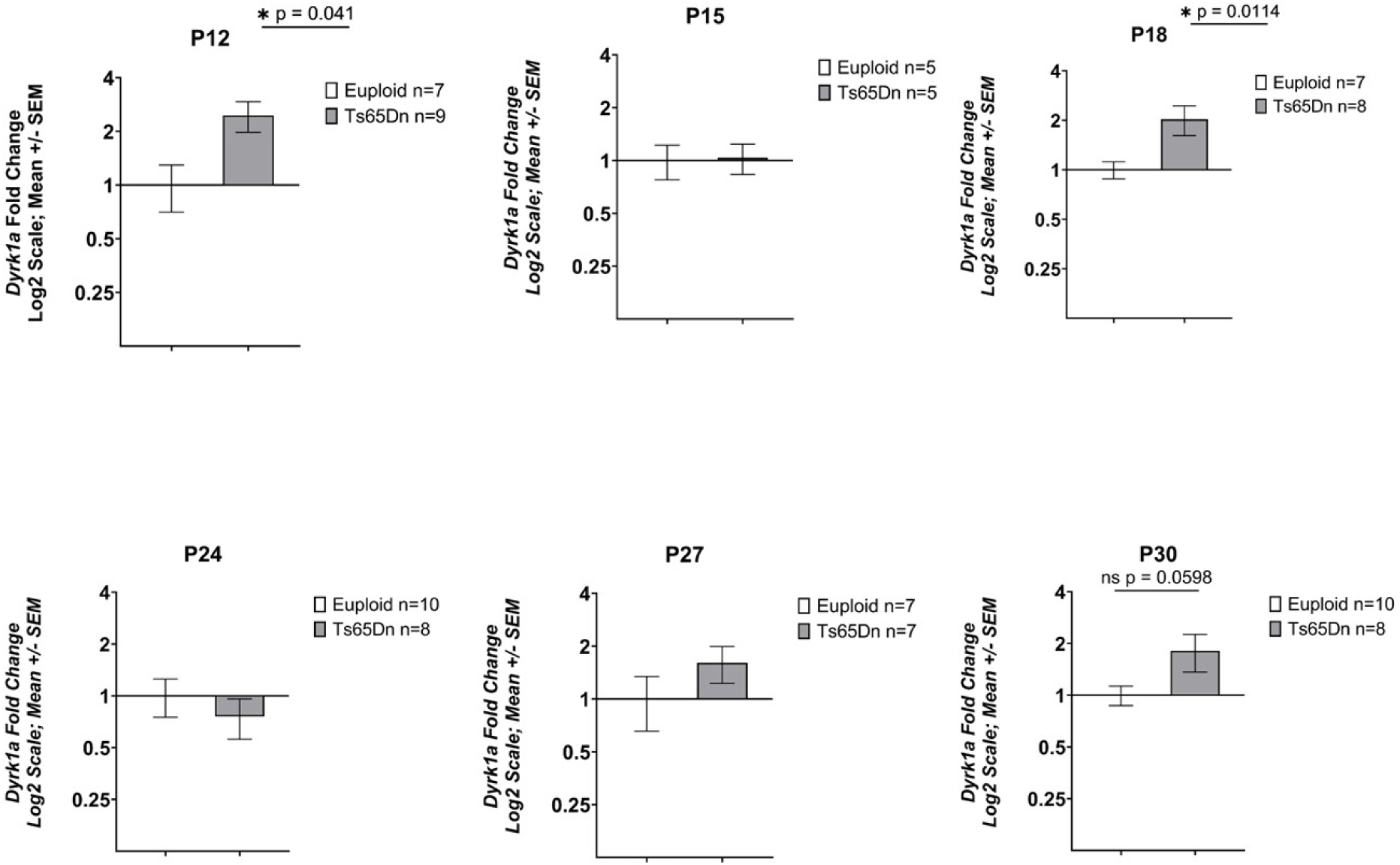
Expression of *Dyrk1a* in Ts65Dn male mice from P12-P42. Data are mean ± SEM. qPCR analysis of cDNA from femoral midshafts represented by fold change on a Log2 scale. Bars and (*) indicate significance value of p < 0.05. P12: euploid (n = 6), Ts65Dn (n = 9); P15: euploid (n = 5), Ts65Dn (n = 5); P18: euploid (n = 7), Ts65Dn (n = 6); P24: euploid (n = 10), Ts65Dn (n = 8); P27: euploid (n = 7), Ts65Dn (n = 7); P30: euploid (n = 10), Ts65Dn (n = 7).

### Treatment of male Ts65Dn mice with CX4945, a putative DYRK1A inhibitor

Given that the germline reduction of *Dyrk1a* in otherwise trisomic male mice resulted in improved trabecular phenotypes at P42 (Blazek et al., 2015a), significant femoral defects emerge at P30 after a period of stagnant development in male Ts65Dn mice, and *Dyrk1a* was trending toward higher expression at P27 and P30, we hypothesized that treatment of male Ts65Dn animals with the DYRK1A inhibitor CX-4945 would improve skeletal formation at P30. Male euploid and Ts65Dn littermates were given 75mg/kg/day CX-4945 suspension (90% PBS:10% DMSO) or vehicle (90% PBS:10% DMSO) via oral gavage from P21 to P29. Mice were euthanized and femurs were collected for μCT analysis on P30.

A repeated-measures ANOVA was performed on body weight data of vehicle and CX-4945 treated euploid and Ts65Dn male mice which showed an effect of age (p<0.001) and an interaction of age and treatment (p=0.014). Pairwise comparisons showed the daily increments in body weight across the 10 days of treatment varied between the vehicle and CX-4945 treated groups, but there were no significant weight differences between vehicle and CX-4945 groups on any treatment day. Two-way ANOVA of average weight from P21 until P30 revealed a genotype effect where Ts65Dn male mice weighed less than euploid mice, which is typical for these mice at this age (Table 2). Weight gained during treatment (weight at P21 subtracted from weight at P30) showed no significant effects of genotype nor treatment, nor their interaction, by two-way ANOVA.

**Table 2:** Significant *p* values from two-way ANOVA (genotype x treatment) for male P30 CX-4945-treated Ts65Dn trabecular and cortical parameters and body weight. All effect and interaction *p* values reported are adjusted using the Benjamini-Hochberg method. ǂ indicates the data was log-transformed for statistical analysis due to non-normal distribution. NS indicates *p* value was not significant. Vehicle-treated euploid (n = 10), CX-4945-treated euploid (n = 13), vehicle-treated Ts65Dn (n = 13), CX-4945-treated Ts65Dn (n = 8).

|  | BMD | BV/TV | Tb.Th | Tb.Sp<sup>‡</sup> | Tb.N |  | Average Weight |  | Weight Gain |
| --- | --- | --- | --- | --- | --- | --- | --- | --- | --- |
| *Genotype Effect* | 0.008 | 0.008 | NS | 0.008 | 0.008 |  | <0.001 |  | NS |
| *Treatment Effect* | NS | NS | NS | NS | NS |  | NS |  | NS |
| *Interaction* | NS | NS | NS | NS | NS |  | NS |  | NS |
|  | Tt.Ar | Ma.Ar | Ct.Ar | Ct.Th | Ps.Pm | Ec.Pm | I<sub>max</sub> | I<sub>min</sub> | Ct.TMD |
| *Genotype Effect* | 0.005 | 0.005 | 0.005 | 0.005 | 0.005 | 0.005 | 0.005 | 0.005 | 0.005 |
| *Treatment Effect* | NS | NS | NS | NS | NS | NS | NS | NS | NS |
| *Interaction* | NS | NS | NS | NS | NS | NS | NS | NS | NS |

Analysis of femoral structure revealed a main effect of genotype in the trabecular BMD (p=0.008), BV/TV (p=0.008), Tb.N (p=0.008) and Tb.Sp (p=0.008) and cortical Tt.Ar (p=0.005), Ma.Ar (p=0.005), Ct.Ar (p=0.005), Ct.Th (p=0.005), Ps.Pm (p=0.005), Ec.Pm (p=0.005), I_max_ (p=0.005), I_min_ (p=0.005), and Ct.TMD (p=0.005) (Table 2). These genotypic differences largely replicated what was previously observed at P30 for Ts65Dn male mice. From these measures, treatment with CX-4945 from P21 to P29 did not improve skeletal defects at P30 in male Ts65Dn mice.

### Temporal reduction of Dyrk1a copy number in maleTs65Dn mice

Given the growth patterns from P24-P30, the altered expression of trisomic *Dyrk1a* in developing mice, normalization of *Dyrk1a* from conception to P42 in Ts65Dn animals restores trabecular and cortical skeletal phenotypes, and CX-4945 treatment did not correct skeletal deficits in male Ts65Dn mice, we hypothesized that the genetic normalization of *Dyrk1a* copy number at P21 would normalize skeletal development of Ts65Dn male mice at P30. To test this hypothesis, Ts65Dn mice were crossed with *Dyrk1a*^fl/wt^ and progeny crossed with a doxycycline-inducible Cre promotor model rtTA^+^, TetOCre^+^. In progeny, doxycycline administration activates Cre expression to excise exon 5 and 6 in one allele of *Dyrk1a* leaving the expression product non-functional (Thompson et al., 2015). Doxycycline administration at weaning (P21) reduced *Dyrk1a* functional copy number in trisomic male Ts65Dn,*Dyrk1a*^fl/wt^, rtTA^+^, TetOCre^+^ (referred to as Ts65Dn,*Dyrk1a*^+/+/dox-cre^) mice during this time.

At P30, there was an effect of genotype in trabecular BMD (p=0.003), BV/TV (p=0.003), Tb.N (p=0.003) and Tb.Sp (p=0.003) and cortical Tt.Ar (p=0.002), Ma.Ar (p=0.002), Ct.Ar (p=0.002), Ct.Th (p=0.002), Ps.Pm (p=0.002), Ec.Pm (p=0.002), I_max_ (p=0.002), I_min_ (p=0.002), and Ct.TMD (p=0.002) (Table 3, Supplemental Table 1). These genotypic differences largely replicated those previously observed between Ts65Dn and euploid littermate male mice at P30 and normalization of *Dyrk1a* copy number, in otherwise trisomic Ts65Dn mice at P21, did not improve skeletal deficits at P30 in male Ts65Dn mice.

**Table 3:**
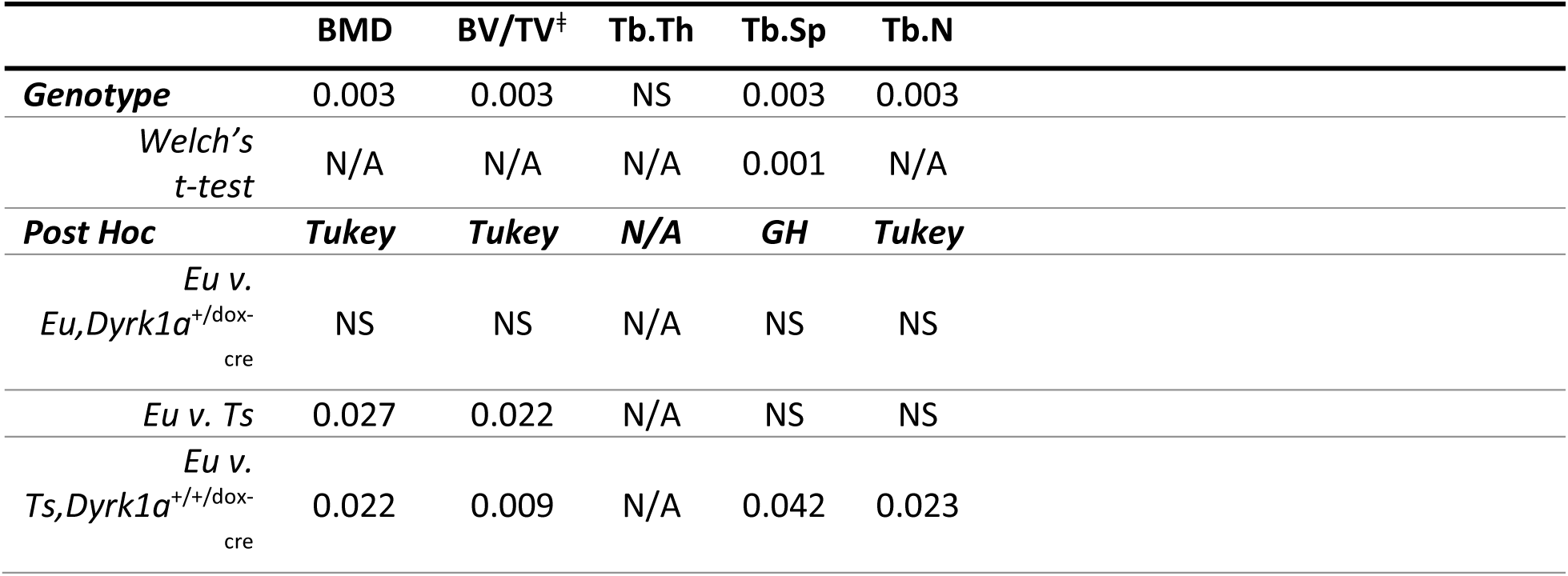
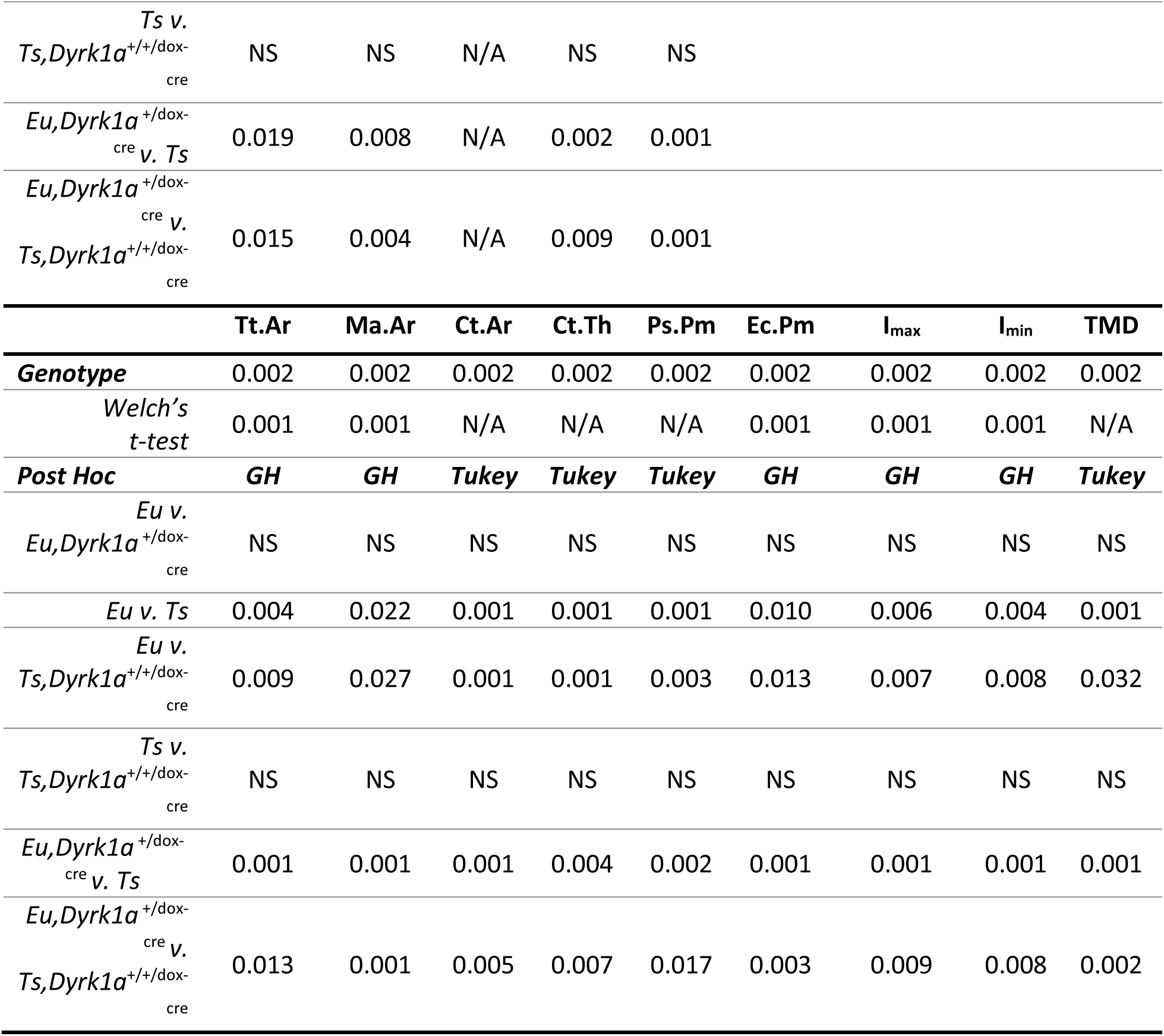
Significant *p* values from one-way ANOVA (genotype) for male P30 Ts65Dn,*Dyrk1a* ^fl/+^,rtTA^+^,TetOCre^+^ (Ts65Dn × *Dyrk1a^+/^*^dox-cre^ offspring) trabecular and cortical parameters. All effect and interaction *p* values reported are adjusted using the Benjamini-Hochberg method. Welch’s t-test was performed to confirm ANOVA results when a significant Levene’s test occurred, indicating unequal variances between groups, and these *p* values are unadjusted. All *p* values from *post hoc* analyses are unadjusted. ǂ indicates the data was log-transformed for statistical analysis due to non-normal distribution. NS indicates *p* value was not significant. N/A indicates Welch’s t-test was not performed because Levene’s test was not significant or *post hoc* analysis was not performed because one-way ANOVA was not significant. GH stands for Games-Howell *post hoc* analysis. Euploid (n = 10), euploid,*Dyrk1a*^+/dox-cre^ (n = 7), Ts65Dn (n = 11), Ts65Dn,*Dyrk1a*^+/+/dox-cre^ (n = 4).

|  | BMD | BV/TV<sup>†</sup> | Tb.Th | Tb.Sp | Tb.N |
| --- | --- | --- | --- | --- | --- |
| **Genotype** | 0.003 | 0.003 | NS | 0.003 | 0.003 |
| *Welch's t-test* | N/A | N/A | N/A | 0.001 | N/A |
| **Post Hoc** | **Tukey** | **Tukey** | **N/A** | **GH** | **Tukey** |
| *Eu v. Eu, Dyrk1a*<sup>+/dox-cre</sup> | NS | NS | N/A | NS | NS |
| *Eu v. Ts* | 0.027 | 0.022 | N/A | NS | NS |
| *Eu v. Ts, Dyrk1a*<sup>+/+/dox-cre</sup> | 0.022 | 0.009 | N/A | 0.042 | 0.023 |

|  |  |  |  |  |  |  |  |  |  |
| --- | --- | --- | --- | --- | --- | --- | --- | --- | --- |
| *Ts v.*<br>*Ts,Dyrk1a*<sup>+/+/dox-cre</sup> | NS | NS | N/A | NS | NS |  |  |  |  |
| *Eu,Dyrk1a*<sup>+/dox-cre</sup><br>*v. Ts* | 0.019 | 0.008 | N/A | 0.002 | 0.001 |  |  |  |  |
| *Eu,Dyrk1a*<sup>+/dox-cre</sup><br>*v.*<br>*Ts,Dyrk1a*<sup>+/+/dox-cre</sup> | 0.015 | 0.004 | N/A | 0.009 | 0.001 |  |  |  |  |
|  | **Tt.Ar** | **Ma.Ar** | **Ct.Ar** | **Ct.Th** | **Ps.Pm** | **Ec.Pm** | **I<sub>max</sub>** | **I<sub>min</sub>** | **TMD** |
| **Genotype** | 0.002 | 0.002 | 0.002 | 0.002 | 0.002 | 0.002 | 0.002 | 0.002 | 0.002 |
| *Welch's*<br>*t-test* | 0.001 | 0.001 | N/A | N/A | N/A | 0.001 | 0.001 | 0.001 | N/A |
| **Post Hoc** | **GH** | **GH** | **Tukey** | **Tukey** | **Tukey** | **GH** | **GH** | **GH** | **Tukey** |
| *Eu v.*<br>*Eu,Dyrk1a*<sup>+/dox-cre</sup> | NS | NS | NS | NS | NS | NS | NS | NS | NS |
| *Eu v. Ts* | 0.004 | 0.022 | 0.001 | 0.001 | 0.001 | 0.010 | 0.006 | 0.004 | 0.001 |
| *Eu v.*<br>*Ts,Dyrk1a*<sup>+/+/dox-cre</sup> | 0.009 | 0.027 | 0.001 | 0.001 | 0.003 | 0.013 | 0.007 | 0.008 | 0.032 |
| *Ts v.*<br>*Ts,Dyrk1a*<sup>+/+/dox-cre</sup> | NS | NS | NS | NS | NS | NS | NS | NS | NS |
| *Eu,Dyrk1a*<sup>+/dox-cre</sup><br>*v. Ts* | 0.001 | 0.001 | 0.001 | 0.004 | 0.002 | 0.001 | 0.001 | 0.001 | 0.001 |
| *Eu,Dyrk1a*<sup>+/dox-cre</sup><br>*v.*<br>*Ts,Dyrk1a*<sup>+/+/dox-cre</sup> | 0.013 | 0.001 | 0.005 | 0.007 | 0.017 | 0.003 | 0.009 | 0.008 | 0.002 |

### Skeletal analysis at P30 of male and female mice with reduction of Dyrk1a copy number from conception

Due to previous observations of significant improvement of trabecular phenotypes in conceptionally reduced Ts65Dn,*Dyrk1a*^+/+/-^ mice (Blazek et al., 2015a), and our current observations with both *Dyrk1a* expression and skeletal development, we hypothesized that a reduction of *Dyrk1a* from conception in otherwise trisomic mice would lead to correction of skeletal deficits in Ts65Dn male mice at P30. Additionally, we wanted to test the hypothesis that female Ts65Dn mice did not have affected trabecular or cortical measures at P30, because female humans with DS exhibit bone deficits later than males (Carfi et al., 2017). To test these hypotheses, femoral skeletal properties were quantified in offspring from (Ts65Dn × *Dyrk1a^+/-^*) matings at P30. There was a significant genotype effect in all trabecular and cortical measures in male mice from a one-way ANOVA: BMD, BV/TV, Tb.Th, Tb.N and Tb.Sp (p=0.002 for all measures) and Tt.Ar, Ma.Ar, Ct.Ar, Ct.Th, Ps.Pm, Ec.Pm, I_max_, I_min_, and Ct.TMD (p=0.002 for all measures) (Figs. 6 and 7). At P30, *post hoc* analyses found that euploid skeletal measures were greater than Ts65Dn, Ts65Dn,*Dyrk1a*^+/+/-^, and euploid,*Dyrk1a*^+/-^ male mice for all trabecular and cortical measures except for Tb.Sp where euploid was less than Ts65Dn and Ts65Dn,*Dyrk1a*^+/+/-^, and cortical Ma.Ar and Ec.Pm, where euploid was greater than Ts65Dn and Ts65Dn,*Dyrk1a*^+/+/-^ mice. A *post hoc* analysis from one-way ANOVA comparison of just Ts65Dn, Ts65Dn,*Dyrk1a*^+/+/-^, and euploid mice showed that euploid mice are significantly different from both trisomic genotypes.

**Figure 6:**
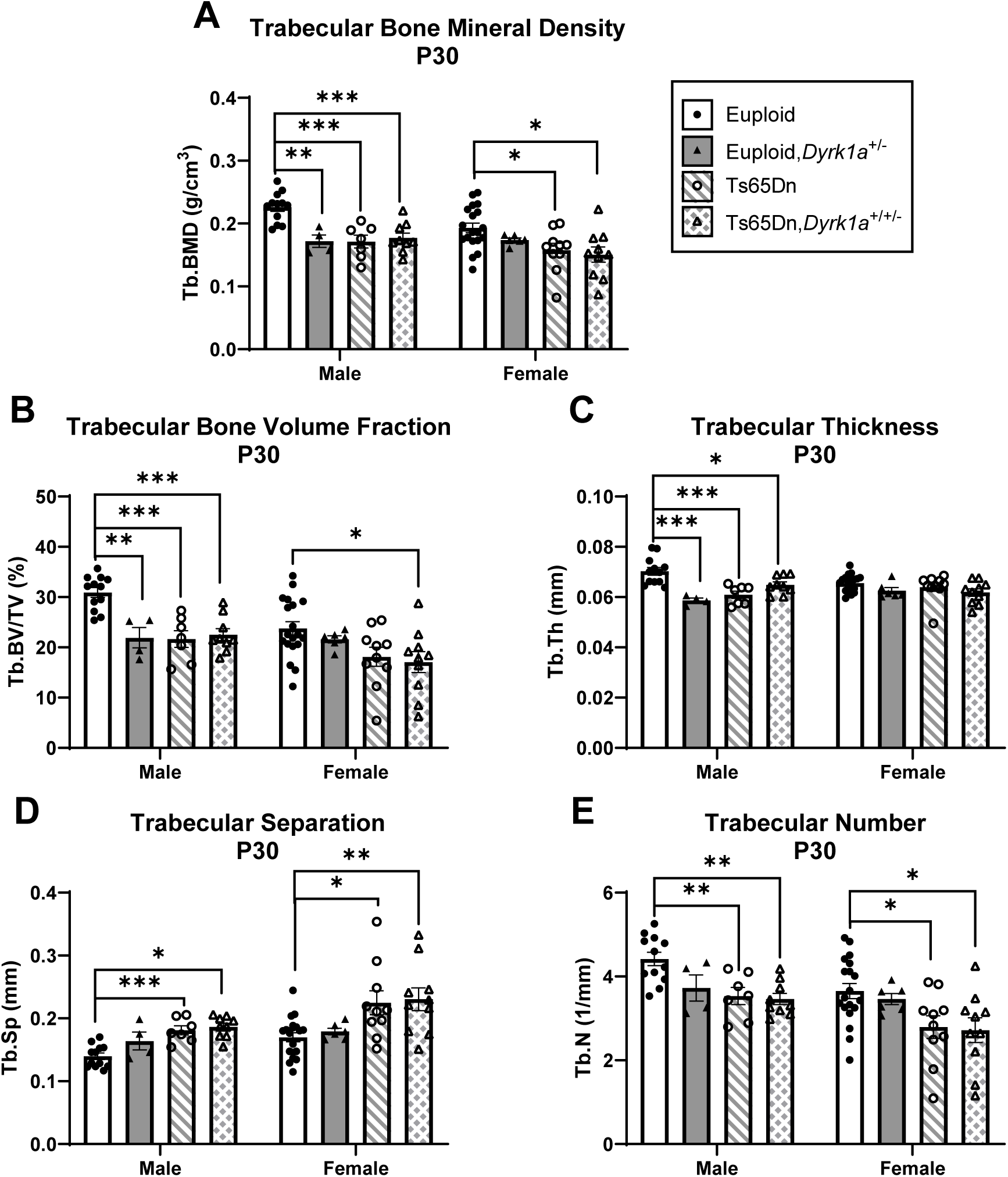
Trabecular analysis of *Dyrk1a* normalization from conception in male and female euploid and Ts65Dn femurs at postnatal day (P)30. Data are mean ± SEM. Male and female mice were analyzed separately using one-way ANOVA and Tukey/Games-Howell *post hoc* analysis. (*) indicates p ≤ 0.05, (**) indicates p ≤ 0.01, (***) indicates p ≤ 0.001. Male: euploid (n = 12), euploid,*Dyrk1a*^+/-^ (n = 4), Ts65Dn (n = 7), Ts65Dn,*Dyrk1a*^+/+/-^ (n = 9); female: euploid (n = 19), euploid,*Dyrk1a*^+/-^ (n = 6), Ts65Dn (n = 10), Ts65Dn,*Dyrk1a*^+/+/-^ (n = 10).

**Figure 7:**
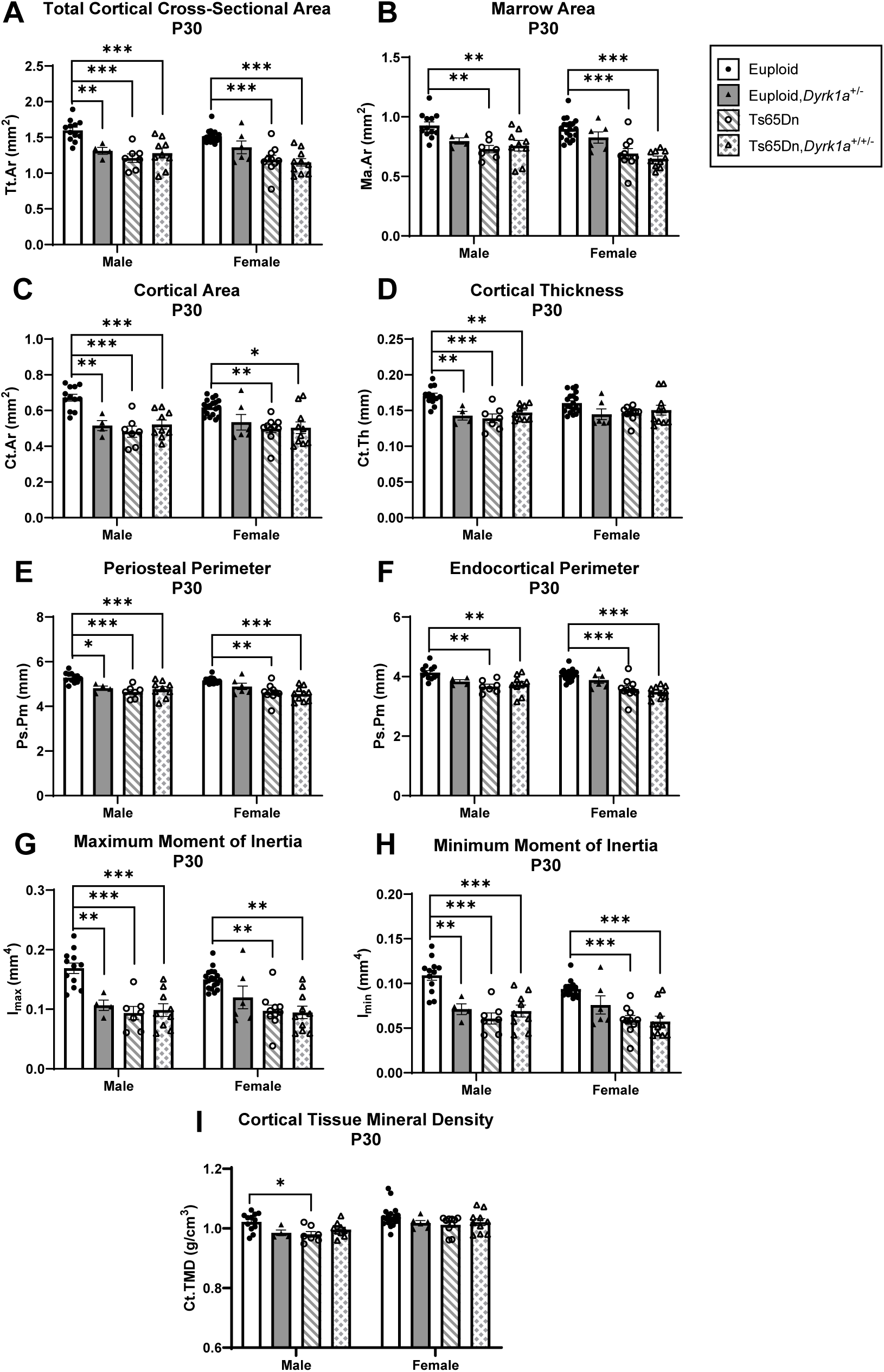
Cortical analysis of *Dyrk1a* normalization from conception in male and female euploid and Ts65Dn femurs at postnatal day (P)30. Data are mean ± SEM. Male and female mice were analyzed separately using one-way ANOVA and Tukey/Games-Howell *post hoc* analysis. (*) indicates p ≤ 0.05, (**) indicates p ≤ 0.01, (***) indicates p ≤ 0.001. Male: euploid (n = 12), euploid,*Dyrk1a*^+/-^ (n = 4), Ts65Dn (n = 7), Ts65Dn,*Dyrk1a*^+/+/-^ (n = 9); female: euploid (n = 19), euploid,*Dyrk1a*^+/-^ (n = 6), Ts65Dn (n = 10), Ts65Dn,*Dyrk1a*^+/+/-^ (n = 10).

For female mice, comparing all offspring from (Ts65Dn × *Dyrk1a^+/-^*) matings at P30 found significant genotype effects in BMD (p=0.018), BV/TV (p=0.020), Tb.N (p=0.015), and Tb.Sp (p=0.005); and Tt.Ar, Ma.Ar, Ct.Ar, Ps.Pm, Ec.Pm, I_max_, and I_min_ (p=0.002 for all measures) (Figs. 6 and 7). *Post hoc* analyses found that for BMD and Tb.N, euploid mice were greater than both Ts65Dn and Ts65Dn,*Dyrk1a*^+/+/-^ mice (Fig. 6A,E). For BV/TV, euploid mice were significantly greater than Ts65Dn,*Dyrk1a*^+/+/-^ mice (Fig. 6B). For Tb.Sp, euploid mice were reduced as compared to Ts65Dn and Ts65Dn,*Dyrk1a*^+/+/-^ mice (Fig. 6D). *Post hoc* analyses for cortical measures found that euploid female mice were greater than both Ts65Dn and Ts65Dn,*Dyrk1a*^+/+/-^ mice (Fig. 7). Taking all data together, Ts65Dn male and female mice exhibit significant deficits in trabecular and cortical bone at P30; these deficits persist even when *Dyrk1a* copy number was returned to normal levels from conception in otherwise trisomic Ts65Dn mice.

### Skeletal analysis at P36 of male and female mice with germline reduction of Dyrk1a copy number

To understand if the skeletal deficits persisted in both male and female Ts65Dn mice and if normalization of *Dyrk1a* copy number would begin to correct these deficits at P36 (halfway between P30 when no correction was seen and P42 where a previous correction was observed), trabecular and cortical skeletal phenotypes were quantified in offspring of (Ts65Dn × *Dyrk1a^+/-^*) matings at P36. For male mice, we found both trabecular [BMD (p=0.005), BV/TV (p=0.006), Tb.Th (p=0.021), Tb.N (p=0.006), and Tb.Sp (p=0.013)] and cortical [Tt.Ar (p=0.004), Ma.Ar (p=0.004),Ct.Ar (p=0.003), Ct.Th (p=0.026), Ps.Pm (p=0.003), Ec.Pm (p=0.004), I_max_ (p=0.003) and I_min_ (p=0.004)] measures were significant by one-way ANOVA (Figs. 8 and 9). In a *post hoc* analysis, euploid male mice were only different from Ts65Dn male mice for BMD, Tb.Th, Tb.N, and Tb.Sp (and euploid,*Dyrk1a*^+/-^ mice for Tb.Th), indicating potential improvement in trabecular bone when *Dyrk1a* copy number was returned to normal in male Ts65Dn mice at P36 (Fig. 8). For cortical deficits, euploid mice were greater than both Ts65Dn and Ts65Dn,*Dyrk1a*^+/+/-^ mice for all measures except Ct.Th, where euploid mice were only significantly greater than Ts65Dn mice, indicating largely no effect of *Dyrk1a* normalization in cortical bone of male Ts65Dn mice at P36 (Fig. 9).

**Figure 8:**
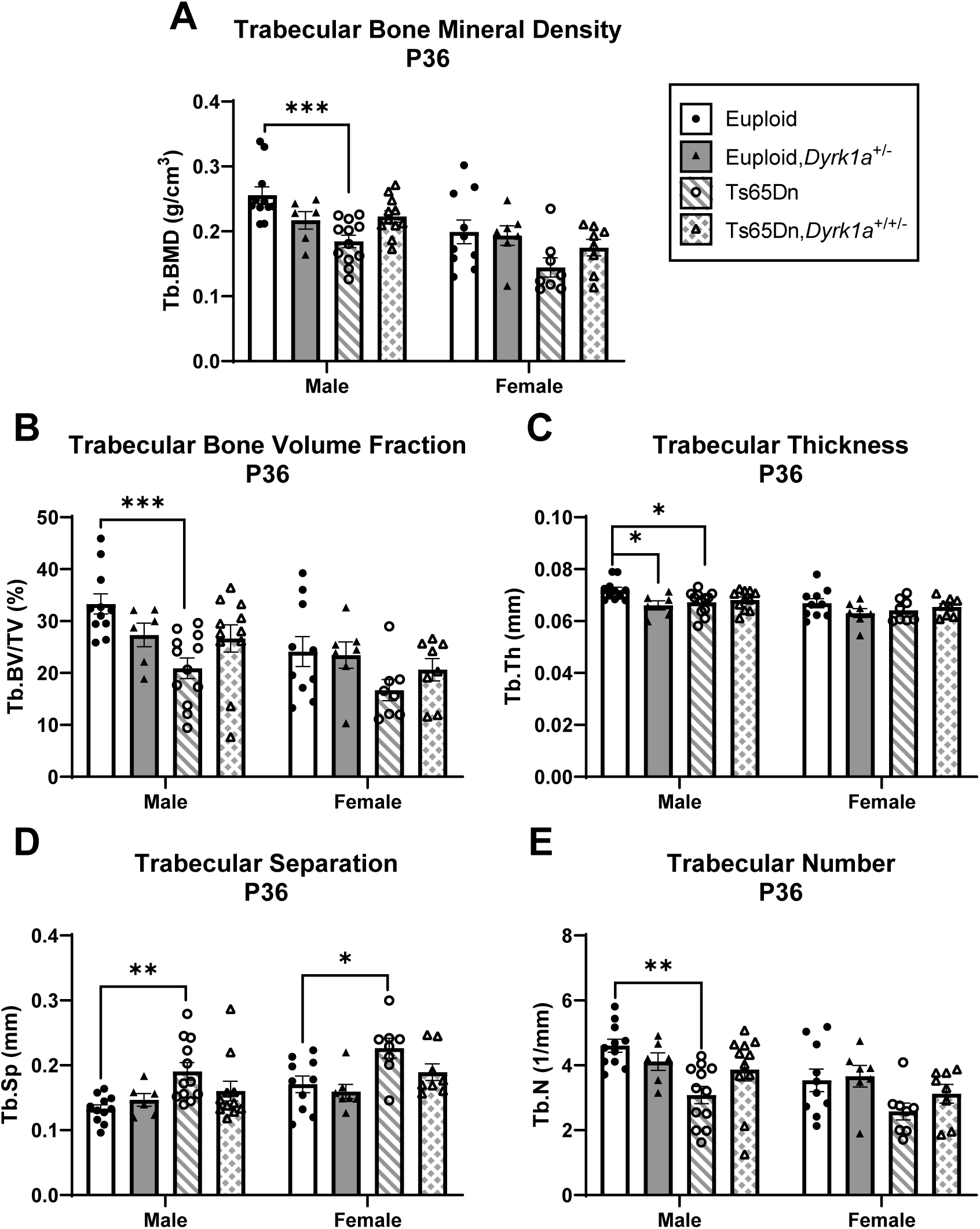
Trabecular analysis of *Dyrk1a* normalization from conception in male and female euploid and Ts65Dn femurs at postnatal day (P)36. Data are mean ± SEM. Male and female mice were analyzed separately using one-way ANOVA and Tukey/Games-Howell *post hoc* analysis. (*) indicates p ≤ 0.05, (**) indicates p ≤ 0.01, (***) indicates p ≤ 0.001. Male: euploid (n = 11), euploid,*Dyrk1a*^+/-^ (n = 6), Ts65Dn (n = 12), Ts65Dn,*Dyrk1a*^+/+/-^ (n = 11); female: euploid (n = 10), euploid,*Dyrk1a*^+/-^ (n = 7), Ts65Dn (n = 8), Ts65Dn,*Dyrk1a*^+/+/-^ (n = 8).

**Figure 9:**
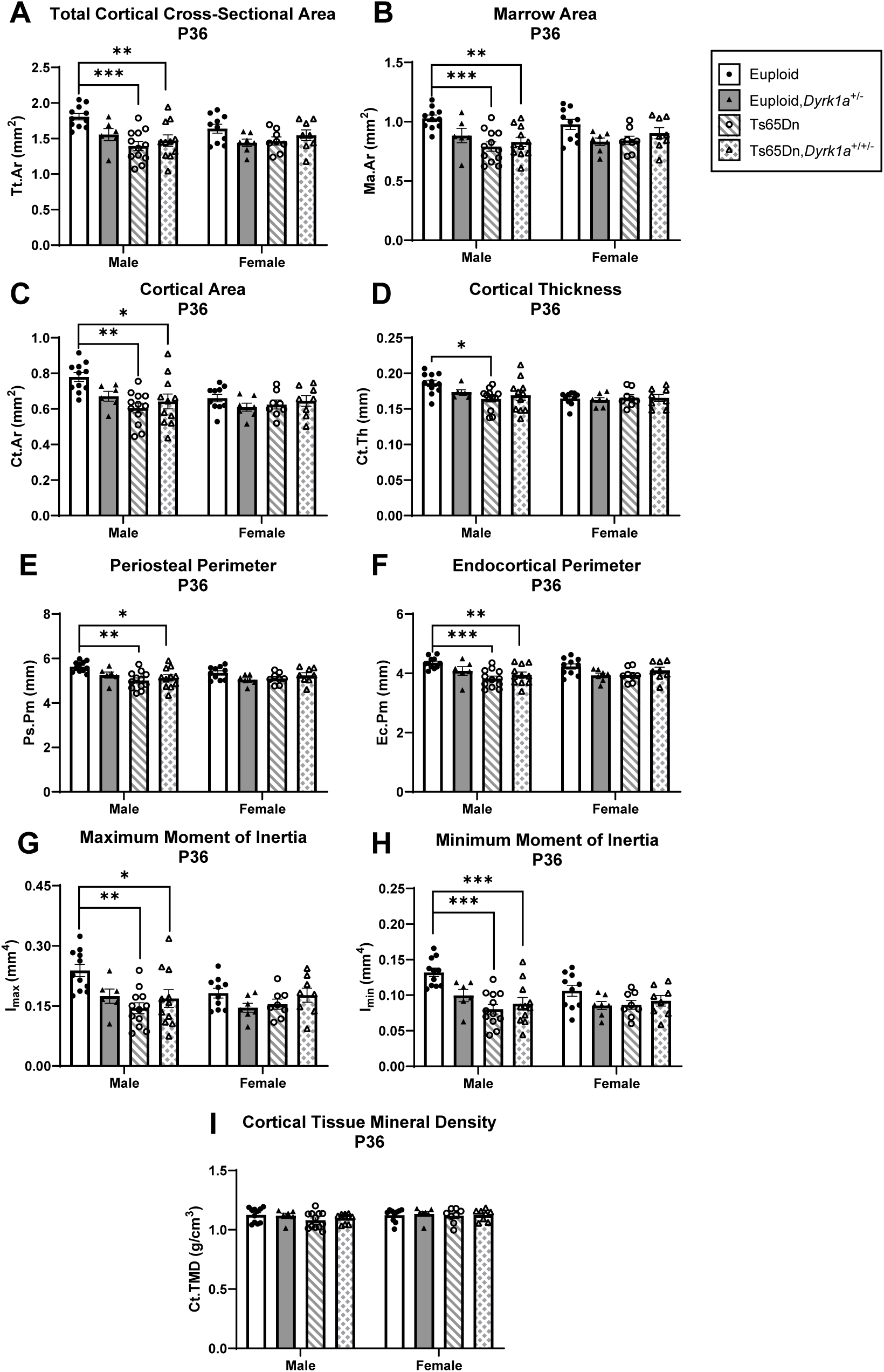
Cortical analysis of *Dyrk1a* normalization from conception in male and female euploid and Ts65Dn femurs at postnatal day (P)36. Data are mean ± SEM. Male and female mice were analyzed separately using one-way ANOVA and Tukey/Games-Howell *post hoc* analysis. (*) indicates p ≤ 0.05, (**) indicates p ≤ 0.01, (***) indicates p ≤ 0.001. Male: euploid (n = 11), euploid,*Dyrk1a*^+/-^ (n = 6), Ts65Dn (n = 12), Ts65Dn,*Dyrk1a*^+/+/-^ (n = 11); female: euploid (n = 10), euploid,*Dyrk1a*^+/-^ (n = 7), Ts65Dn (n = 8), Ts65Dn,*Dyrk1a*^+/+/-^ (n = 8).

For female offspring of (Ts65Dn × *Dyrk1a^+/-^*) matings at P36, the only difference observed was Tb.Sp where euploid and euploid,*Dyrk1a*^+/-^ mice had significantly less trabecular separation than Ts65Dn and Ts65Dn,*Dyrk1a*^+/+/-^ mice (Fig. 8D). No other trabecular or cortical parameters were significantly different in female offspring of (Ts65Dn × *Dyrk1a^+/-^*) matings at P36 (Figs. 8 and 9).

To better understand how other influences besides *Dyrk1a* copy number were affecting trisomic bone development from P24-P30, during the stagnant growth that results in lasting defects, we examined mRNA expression in femurs from genes that have been linked to skeletal development: *Bglap*, *Runx2*, *Rbl2*, and *Alpl* at P24, P27, and P30 (Rodda and McMahon, 2006, Gao et al., 2014). *Rbl2*, also known as *P130*, is a gene transcribed via the transcription factor FOXO1 (Kops et al., 2002). Nuclear FOXO1 competes with TCF/LEF1 for β-catenin to transcribe target genes but is also exported from the nucleus after phosphorylation by DYRK1A (Bhansali et al., 2021). *P130* is hypothesized to indirectly report DYRK1A activity (Forristal et al., 2014); increased DYRK1A activity would result in reduced expression of *Rbl2*. In trisomic male Ts65Dn mice at P24, P27, and P30, *Bglap*, *Runx2*, *Rbl2*, and *Alpl* levels were not statistically different than euploid littermate mice (Supplemental Figure 2).

## Discussion

### Initiation of bone deficits in male Ts65Dn mice

Male individuals with DS have bone deficits at earlier ages then female individuals with DS as compared to those without DS. Skeletal deficits in Ts65Dn mice have shown a similar trend, with persistent skeletal deficits occurring in male trisomic mice affected earlier than female mice. This study of male Ts65Dn mice during postnatal development shows that cortical deficits are present in male trisomic as compared to euploid mice beginning at P12, and some trabecular deficits begin to present at P18. Some trisomic trabecular and cortical measurements are near equivalent to euploid male mice at P24 and seem to exhibit stagnant development in male Ts65Dn mice until P30, when male trisomic trabecular and cortical deficits are fully present again. In male Ts65Dn mice, trabecular and cortical deficits persist at P36, and as we have previously shown, at P42 (6 weeks) and 16 weeks. Others have noted similar trabecular and cortical deficits in tibiae of male Ts65Dn mice at 8 weeks and 24 months (Fowler et al., 2012, Williams et al., 2018). Taken together, in male Ts65Dn mice, cortical bone deficits are present at most every developmental age tested, whereas trabecular deficits are more transient, but become significantly deficient during development. The oscillations in trabecular phenotypes in male mice might be due to altered expression of *Dyrk1a* and/or other trisomic genes during certain time points based on our qPCR analysis of bone. It is unknown if a similar pattern of oscillating bone phenotypes is seen in humans with DS because longitudinal long bone studies have not been completed in humans with DS. This could partially explain inconsistencies between studies of bone phenotypes in adolescents with DS.

### Differences in skeletal deficits between trisomic male and female mice

In previous findings, we demonstrated that at P42, female Ts65Dn mice showed inferior femoral trabecular properties in BMD and Tb.Sp, and also showed reduced cortical measures including Tt.Ar, Ma.Ar, Ps.Pm, and Ec.Pm as compared to female euploid mice (Thomas et al., 2021). At P30, we observed that female trisomic mice from a (Ts65Dn x *Dyrk1a*^+/-^) cross had deficits in femoral trabecular BMD, Tb.N, and Tb.Sp and most cortical skeletal measures as compared to euploid mice. At P36, however, femurs from female trisomic mice only had increased Tb.Sp and no significant cortical deficits. These data indicate that female trisomic mice have some appendicular bone deficits at P30, followed by a period of near normal bone characteristics at P36 that becomes significantly worse by P42. Taken together, both trisomic male and female mice have periods of significant femoral deficits, periods of recovery to normal measures in a sex-specific manner (around P24 for males and P30 for females), then become worse over time, with male Ts65Dn mice showing consistent bone deficits earlier (around P30) than female mice (between P36 and P42).

In Dp1Tyb mice, a duplication mouse model of DS, trabecular deficits also are observed in male Dp1Tyb mice at P42 before female Dp1Tyb mice, consistent with Ts65Dn mice; however, cortical deficits are in both sexes at all ages investigated, which is inconsistent with Ts65Dn mice, but this may be due to the limited number of timepoints investigating both sexes of Dp1Tyb mice. Female Dp1Tyb mice are also discordant from female Ts65Dn mice in that they have not been found to have trabecular deficits, even at 16 weeks.

### Relationship of trisomic Dyrk1a to bone deficits in Ts65Dn mice

Three copies of *Dyrk1a* have been hypothesized to contribute to bone deficits and these data further clarify the timing of trisomic *Dyrk1a* involvement appendicular skeletal deficits in Ts65Dn mice. Trisomic *Dyrk1a* was not consistently overexpressed in femoral RNA of male mice at all timepoints from P12-P30. We have shown in brains of Ts65Dn mice, DYRK1A protein and mRNA are not always consistently overexpressed in cerebellum, hippocampus, and cerebral cortex throughout development (Hawley et al., 2023). Although *Dyrk1a* RNA is significantly overexpressed in the femurs of male Ts65Dn as compared to control mice at P12 and P18 and trending to higher levels at P27 and P30, the normalization of *Dyrk1a* copy number in otherwise trisomic male mice from P21 to P30, or from conception, did not have a corrective effect on male or female trisomic trabecular or cortical femoral phenotypes at P30. At P36, there was no effect of trisomy or normalization of *Dyrk1a* copy number from conception in otherwise trisomic female mice, but in Ts65Dn,*Dyrk1a*^+/+/-^ male mice, trabecular deficits and Ct.Th were improved. Previous data from our lab showed that not all appendicular skeletal measures were affected by normalizing *Dyrk1a* expression, cortical measures in particular (Blazek et al., 2015a, Sloan et al., 2023). Similarly, there is not an apparent connection between oscillating *Dyrk1a* expression and cortical femoral phenotypes in perinatal development. Taken together, these data indicate that 1) trisomic *Dyrk1a* expression changes throughout perinatal development in the femur, challenging the dogma in the field that all genes are consistently upregulated 1.5-fold in trisomic mice at all times; 2) trisomic *Dyrk1a* begins to have a significant effect on trabecular and some cortical femoral phenotypes in Ts65Dn male mice at P36 even though there are ages where *Dyrk1a* is overexpressed or is trending overexpression in this bone; and 3) femoral bone is likely affected in female Ts65Dn mice after P36 but before P42, yet the effect of trisomic *Dyrk1a* on female trisomic femurs is still unknown. How trisomic *Dyrk1a* affects skeletal phenotypes may change as Ts65Dn mice increase in age, and the interaction between other trisomic genes may affect the developing trabecular and cortical deficits in at least male Ts65Dn mice (Sloan et al., 2023).

### CX-4945 treatment in male Ts65Dn mice

Treatment with CX-4945 tested the hypothesis that treatment using a putative DYRK1A inhibitor during a window of *Dyrk1a* overexpression and stagnating trisomic femoral development would improve femoral bone in Ts65Dn mice. CX-4945 was not soluble in water or PBS, limiting initial dissolution to DMSO. Due to health concerns over multiple treatments with DMSO, the total percentage of each treatment in this study was limited to 10% total DMSO. CX-4945 did not remain dissolved at 90% PBS:10% DMSO, however, it formed a suspension that was relatively stable at 37°C. Treatment of male Ts65Dn mice from P21-P29 with 75mg/kg/day CX-4945 in a 90% PBS:10% DMSO suspension via oral gavage proved ineffective in improving *Dyrk1a*-related femoral phenotypes. This lack of improvement in the femur could be because of the limited effect of CX-4945 as a DYRK1A inhibitor in bone, or because treatment was given when overexpression of *Dyrk1a* did not seem to affect bone development.

More unexpected was the effect of vehicle DMSO treatment on Ts65Dn mice; the mean BV/TV in vehicle treated mice was approximately 23%, compared to untreated P30 Ts65Dn mice characterized earlier which had a mean approximate BV/TV of 15% (Supplemental table 2). Vehicle treatment of male Ts65Dn mice from P21 to P29 appeared to improve trabecular measures to euploid levels. However, not all *Dyrk1a*-related femoral phenotypes were affected; Ct.Th was not significantly different between vehicle-treated and untreated Ts65Dn mice, suggesting a positive effect only on trabecular bone. Previous reports administering DYRK1A inhibitors via oral gavage showed significant differences in *Dyrk1a*-related measures in Ts65Dn mice receiving the vehicle (Goodlett et al., 2020, Stringer et al., 2017a), suggesting that 10% DMSO, as part of a vehicle treatment, improved trabecular defects.

DMSO has been reported to decrease osteoclast maturation in a dose-dependent manner (Lemieux et al., 2011, Yang et al., 2015). Others have reported increased osteoblast differentiation and activity in mouse and human osteoblasts treated with DMSO, along with increased osteogenic gene expression (Stephens et al., 2011, Cheung et al., 2006). It may be that DMSO interacts with products of trisomy to significantly increase femoral measures in Ts65Dn, but not euploid mice. While unlikely, it is possible that because the CX-4945-treated cohorts were treated before vehicle-treated cohorts, seasonal growth effects may have impacted analysis between the two groups. It is possible that nutritional aspects may play a role; mean body weight was increased at P30 in vehicle-treated Ts65Dn mice (15.72 g) compared to untreated Ts65Dn mice (12.74 g) but no difference between vehicle-treated (18.58 g) and untreated (19.63 g) euploid mice. This illustrates the importance of using vehicle treatment in therapeutic studies on DS and other models.

### Implications for treatments of bone deficits

Overexpression of *Dyrk1a* during crucial developmental stages likely contributes to DS-related phenotypes such as cognitive impairment, skeletal deficits, and craniofacial abnormalities (Hawley et al., 2023, Stringer et al., 2017b, McElyea et al., 2016). While interventions targeting *Dyrk1a* overexpression using genetic and therapeutic mechanisms have shown limited normalization of some skeletal and neurodevelopmental deficits, the potential efficacy of translational DYRK1A inhibitors is still questionable (Stringer et al., 2017b). We have shown expression of trisomic *Dyrk1a* varies spatially and temporally in bone and brain and may affect a wide range of DS-associated phenotypes. Because of the dynamic nature of *Dyrk1a* overexpression and the dosage effects of both over- and under-expression of *Dyrk1a*, care must be taken when administering DYRK1A inhibitors. Mutations in *Dyrk1a* lead to Dyrk1a syndrome that includes both neurological and skeletal phenotypes (Ji et al., 2015, Huang et al., 2023). A general pharmacological reduction of DYRK1A, especially during development, may have detrimental effects in tissues where *Dyrk1a* is not overexpressed. Administration of a DYRK1A inhibitor during an incorrect window of time may not improve the targeted deficit, whereas a later administration of the inhibitor may have positive effects. A negative result of an inhibitor may not be due to the limited efficacy or off-target effects of a potential DYRK1A inhibitor, but rather to incorrect timing and tissue specificity of the inhibitor.

Additionally, differences in expression of trisomic *Dyrk1a* between the sexes must be accounted for when determining therapeutic approaches. Differential expression of DYRK1A protein and RNA between male and female Ts65Dn mice has been shown in the cerebellum, cerebral cortex, and hippocampus (Hawley et al., 2022, Hawley et al., 2023), and the differential effects of trisomic *Dyrk1a* on femur phenotypes is now apparent (this study). From these studies of trisomic Dyrk1a expression, it appears that treatments to inhibit DYRK1A would need to differ between the sexes because of the spatiotemporal differences in *Dyrk1a* overexpression. From these studies, attempted normalization of DYRK1A expression would need to begin in males (before P36) before females and continue through the major influences of trisomic DYRK1A on bone formation to correct femoral deficits.

### Study Limitations

There are many DS mouse models in which bone deficits have been characterized, including Ts65Dn, Dp1Tyb, Dp1Rhr, and Dp(16)1Yey (Blazek et al., 2011, Thomas et al., 2020, Sherman et al., 2022, Sloan et al., 2023). Ts65Dn mice are the most well characterized DS mouse model and appear to effectively model femoral DS-associated skeletal deficits, but skeletal studies in female mice have been limited because of the need to utilize these mice to generate additional mice. Furthermore, Ts65Dn mice also contain triplication of ∼35 protein coding genes that are not orthologous to Hsa21. The recently generated Ts66Yah mouse removes these non-orthologous genes (Duchon et al., 2022), and skeletal abnormalities in these mice have not yet been characterized. Additionally, offspring used to identify the influence of trisomic *Dyrk1a* in study of *Dyrk1a* skeletal deficits from P12-P30 came from (Ts65Dn × B6C3.*Dyrk1a^fl/fl^*) matings. Ts65Dn and euploid mice should be equivalent on the ∼50% B6 and ∼50% C3H background from these studies, but there may have been intrauterine influences from male mice used in these crosses.

Due to the composition of the bones at these ages, functional analysis (3-point mechanical bending) could not be performed so the direct strength of these bones was not measured. However, maximum moment of inertia (I_max_) and minimum moment of inertia (I_min_), calculated from cortical cross-sections, can be used as an indication of the bone’s response to force and to estimate bone strength (Cole and van der Meulen, 2011, Wallace, 2019)

## Conclusion

This study identifies key timepoints in Ts65Dn DS mouse model appendicular skeletal development for targeted treatment of skeletal deficits associated with triplicated *Dyrk1a*. Male Ts65Dn mice have cortical deficits earlier (P12) than trabecular deficits, which are not consistently altered until P30. Periodic normalization of femoral phenotypes was also identified in male and female Ts65Dn mice. Despite having triplicated *Dyrk1a*, *Dyrk1a* mRNA levels are not always overexpressed in male Ts65Dn femurs. *Dyrk1a* normalization improves skeletal phenotypes in male mice after P30, and this effect is largely limited to trabecular bone. This indicates a complex relationship between *Dyrk1a* expression, and skeletal phenotypes associated with Ts21 that may be influenced by several factors including age, sex, and timing of *Dyrk1a* overexpression. Our data from female Ts65Dn mice suggest consistent trabecular and cortical deficits may not arise until P42 (after male Ts65Dn mice), but later ages need to be analyzed to confirm this. Together, this illustrates the importance of identifying spatial and temporal windows of development and gene expression in DS mouse models to improve pre-clinical treatment outcomes.

## Competing Interests

The authors have no competing interests to declare.

## Funding

This research in this manuscript was supported by funds from National Institutes of Health Eunice Kennedy Shriver National Institute for Child Health and Human Development HD090603 and the National Institute of Arthritis and Musculoskeletal and Skin Diseases AR078663 (RJR). Additional funding was provided by Undergraduate Research Opportunities Program grant from the Center for Research and Learning at IUPUI (MPB and IC)

## CRediT Authorship Contribution Statement

Jonathan M. LaCombe: Conceptualization, Formal analysis, Investigation, Visualization, Writing – original draft preparation, Project administration

Kourtney Sloan: Formal analysis, Visualization, Writing – review and editing

Jared R. Thomas: Investigation, Formal analysis, Visualization, Writing – review and editing

Matthew P. Blackwell: Investigation, Formal analysis, Writing – review and editing, Funding acquisition

Isabella Crawford: Investigation, Formal analysis, Writing – review and editing, Funding acquisition

Joseph M. Wallace: Software, Resources, Writing – review and editing

Randall J. Roper: Conceptualization, Resources, Writing – original draft preparation, Funding acquisition, Supervision, Project administration

## Supplemental Figures

**Supplemental Figure 1.**
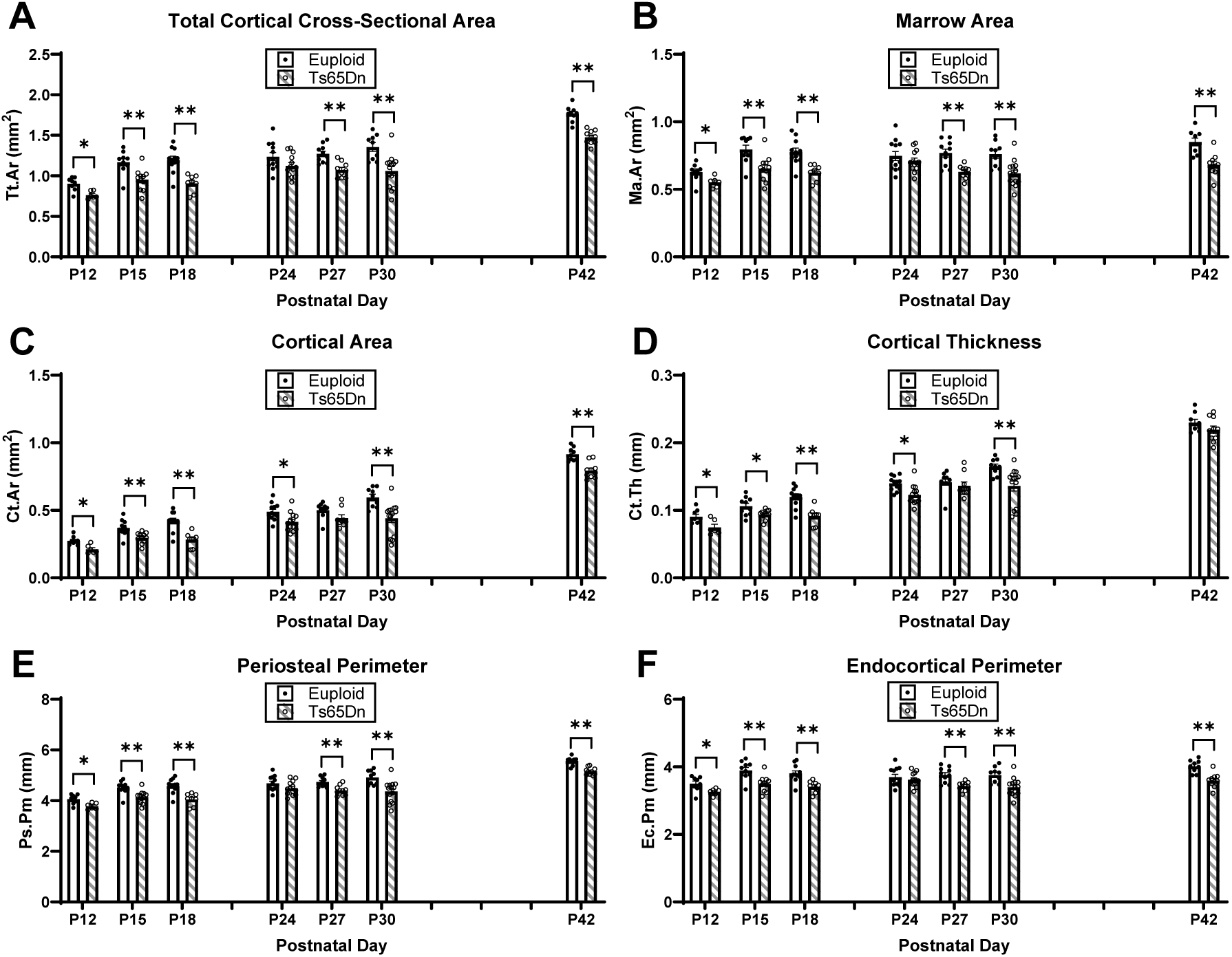
Cortical analysis of male Ts65Dn femurs from postnatal day (P)12-42. Remaining cortical parameters from μCT analysis. Data are mean ± SEM. Significance determined through two-tail t-test with FDR adjustment. (*) indicates p ≤ 0.05, (**) indicates p ≤ 0.01.

**Supplemental Figure 2.**
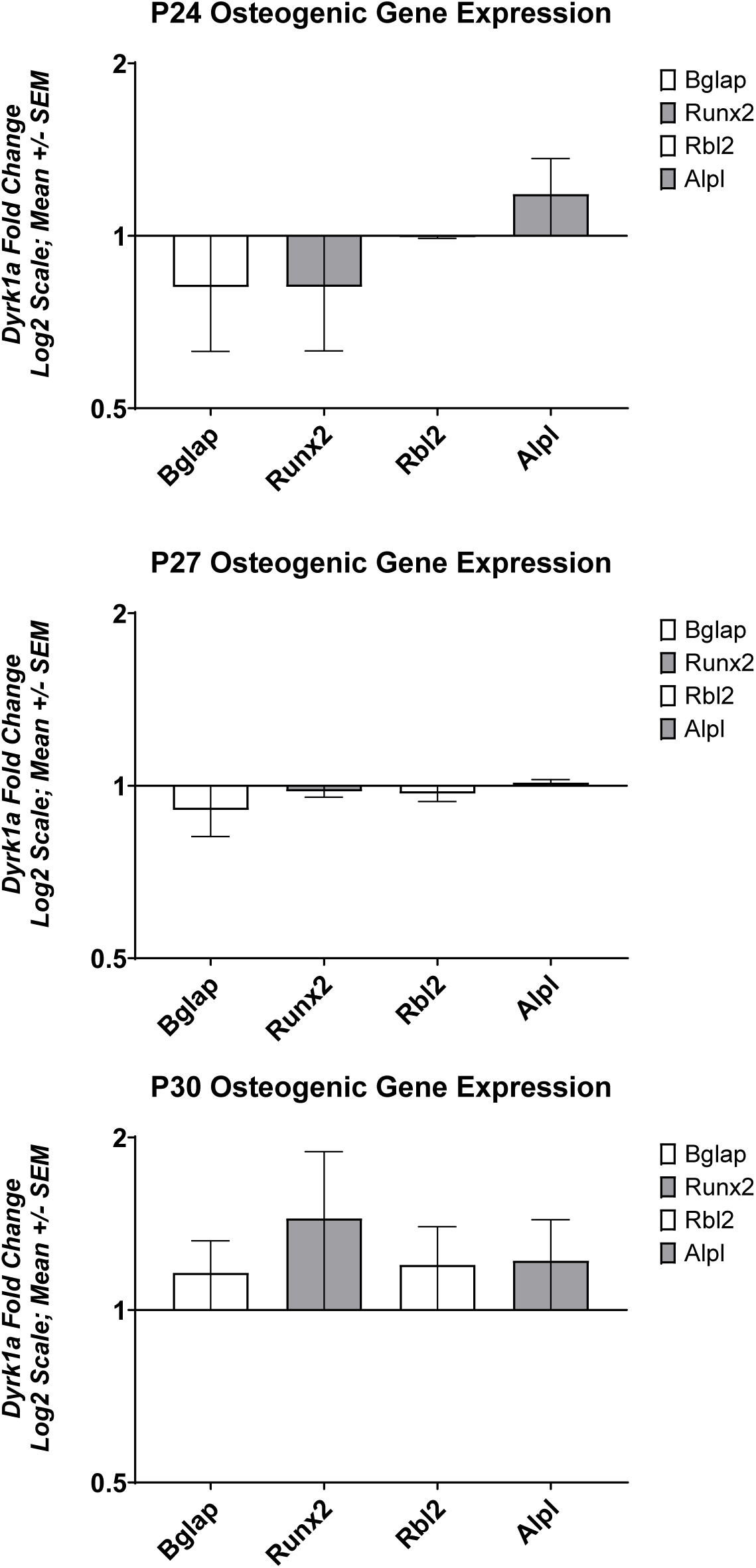
Osteogenic gene expression at P24, P27, and P30

## Supplemental Tables

**Supplemental Table 1.**
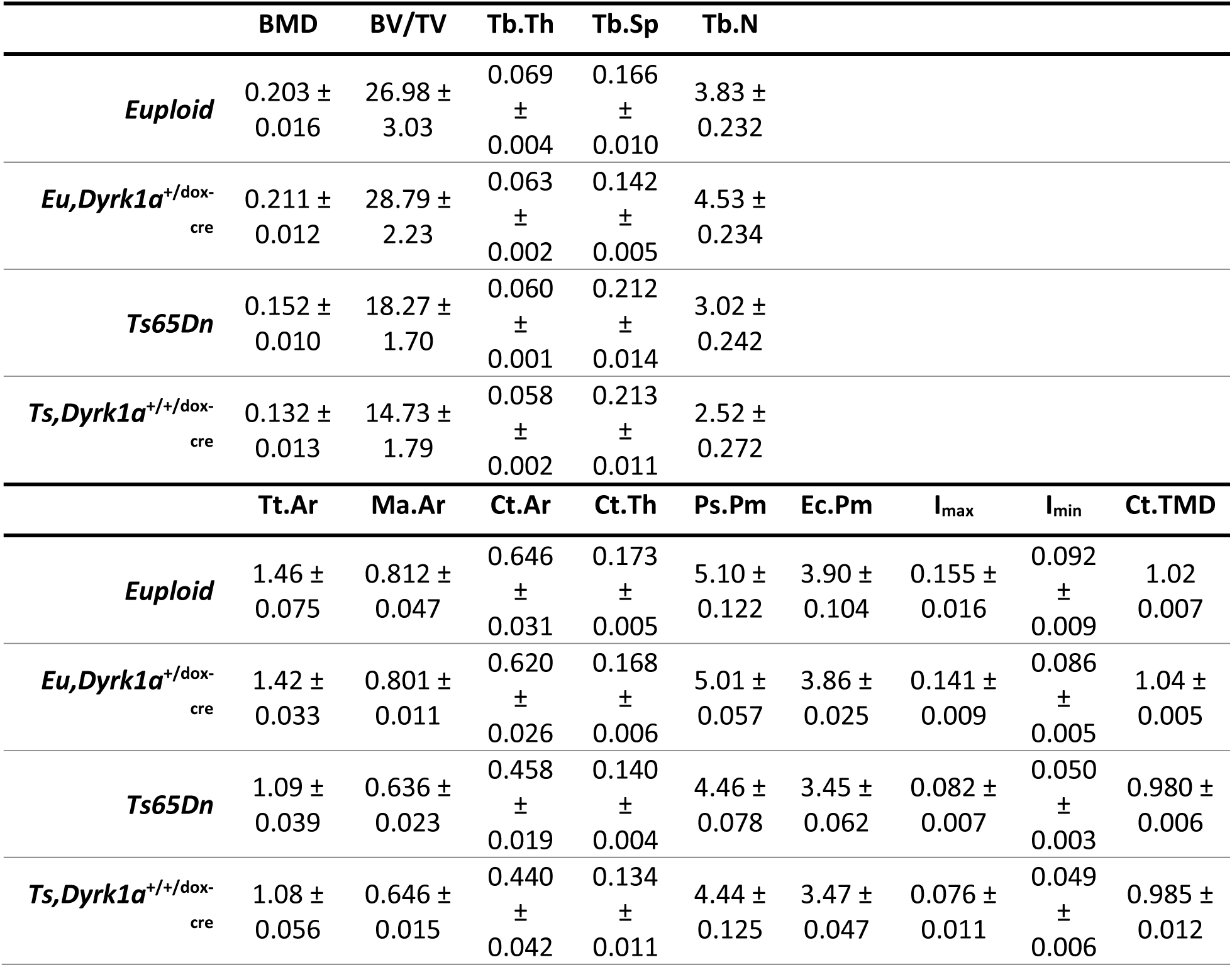
Average and standard error of the mean (SEM) of µCT parameters for temporal reduction of *Dyrk1a* copy number in maleTs65Dn mice at P30.

**Supplemental Table 2.**
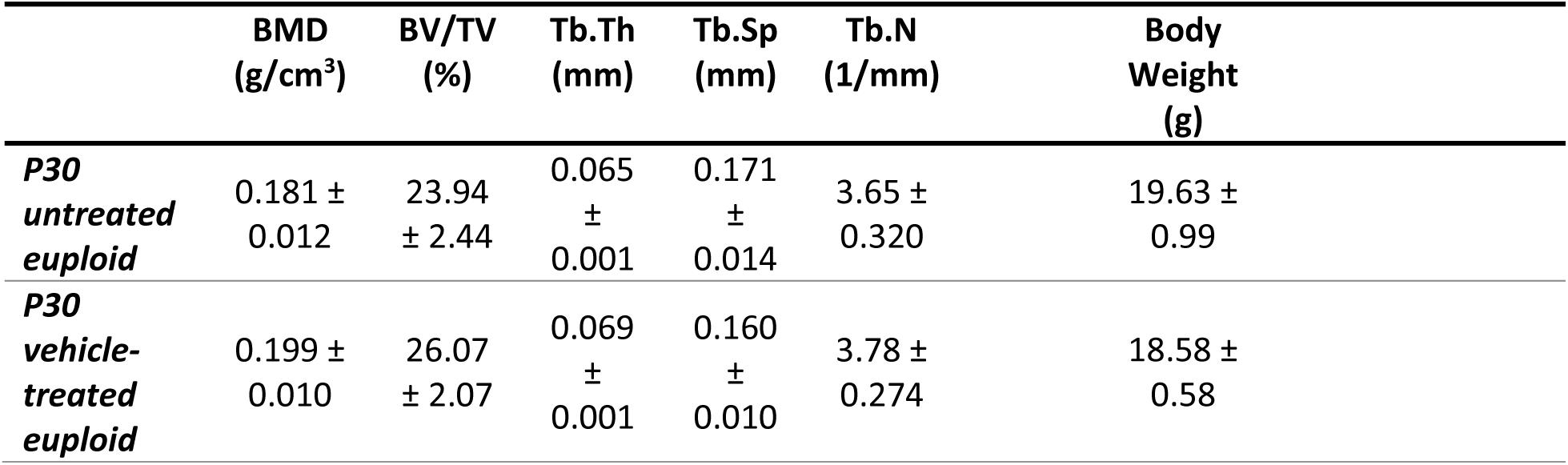
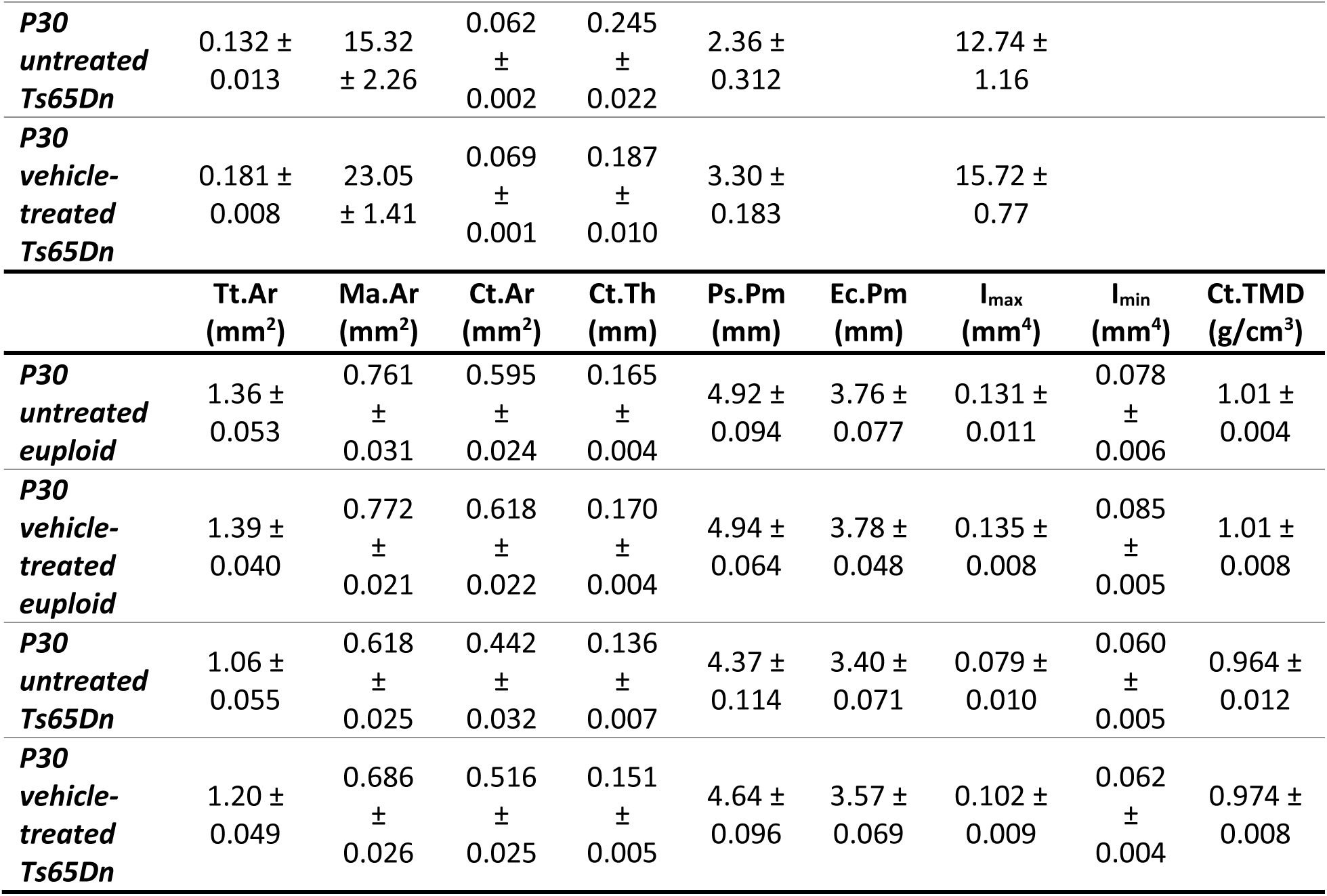
Comparison of P30 vehicle (10%DMSO:90%PBS)-treated and untreated euploid and Ts65Dn mice. Data are average ± SEM.

## Notes

### Competing Interest Statement

The authors have declared no competing interest.

